# Thousands of human mutation clusters are explained by short-range template switching

**DOI:** 10.1101/2021.11.26.470150

**Authors:** Ari Löytynoja

## Abstract

Variation within human genomes is unevenly distributed, and variants show spatial clustering. DNA-replication-related template switching is a poorly known mutational mechanism capable of causing major chromosomal rearrangements as well as creating short inverted sequence copies that appear as local mutation clusters in sequence comparisons. I reanalyzed haplotype-resolved genome assemblies representing 25 human populations and multinucleotide variants aggregated from 140,000 human sequencing experiments. Local template switching could explain thousands of complex mutation clusters across the human genome, the loci segregating within and between populations. I developed computational tools for identification of template switch events using both short-read sequencing data and genotype data, and for genotyping candidate loci using short-read data. The characteristics of template-switch mutations complicate their detection and, worryingly, widely used analysis pipelines for short-read sequencing data, normally capable of identifying single nucleotide changes, were found to miss template-switch mutations of tens of base pairs, potentially invalidating medical genetic studies searching for a causative allele behind genetic diseases. Combined with the massive sequencing data now available for humans, the novel tools described here enable building catalogs of affected loci and studying the cellular mechanisms behind template switching in both healthy organisms and disease.

## Introduction

Twenty years after the publication of the draft genomes (Venter et al. 2001; Lander et al. 2001), the Telomere-to-Telomere Consortium reported the first truly complete sequence of a human genome (Nurk et al. 2022; Aganezov et al. 2022). However, every genome is unique, and the challenge now is to understand the global genomic diversity in the human population (Karczewski et al. 2020; Taliun et al. 2021) and to build a comprehensive pangenome to represent this variation (Miga and Wang 2021). While single nucleotide variation (SNV) and short insertion-deletions (indels) have been resolved relatively accurately with classical short-read sequencing (Bentley et al. 2008), the characterization of structural variants (SVs) and low-complexity sequences has been lacking. The Human Genome Structural Variation Consortium developed a method (Porubsky et al. 2021) for phased diploid genome assembly with combination of long-read PacBio whole-genome sequencing (WGS; Eid et al. 2009) and Strand-seq phasing (Falconer and Lansdorp 2013) data. Ebert et al. (2021) applied this method to 32 diverse human individuals and produced 64 assembled haplotypes, i.e., maternal and paternal copies of the genome. With the help of phased diploid genomes, they massively expanded the catalog of known SVs and, having the information of coinherited nearby sequence differences, could study in detail the different subclasses of complex variants. Such accurate characterization of multiple genomes will lay the foundations for understanding the role of complex mutations in human phenotypes and disease (Zhang et al. 2009; Weischenfeldt et al. 2013; Sakamoto et al. 2020) and in evolution (Zhang et al. 2009; Perry et al. 2007; Yan et al. 2021). More generally, the unprecedented resolution of the new genome data will open novel possibilities for understanding the mutational mechanisms of the complex eukaryotic genomes (Conrad et al. 2011; Veltman and Brunner 2012; Śegurel et al. 2014).

Although many subclasses of variants studied by Ebert et al. are clearly independent, mutations are known to be clustered (Śegurel et al. 2014) and their separation at the analysis stage may dismiss crucial information. We compared earlier two independent assemblies of the human genome (International Human Genome Sequencing Consortium 2004; Levy et al. 2007) and found mutation clusters consistent with DNA-replication related template switching (Löoytynoja and Goldman 2017). The mechanism, originally described and studied in bacteria (Ripley 1982; Dutra and Lovett 2006), resembles FoSTeS (Fork Stalling and Template Switching; Lee et al. 2007) and MMBIR (Microhomology-Mediated Break-Induced Replication; Hastings et al. 2009) but is local and reciprocal switches typically occur within a region of a few tens or hundreds of base pairs (bp). Unlike the chromosomal rearrangements created by FoSTeS and MMBIR, the footprint of local template switch mutations (TSMs) are short inversions, either happening in place or creating a reverse complement copy of a nearby sequence region; these are believed to be produced by the replication briefly switching either to the complementary DNA strand or going backwards along the nascent DNA strand (Ripley 1982; Dutra and Lovett 2006) (Fig. S1). In sequence comparisons, mutations compatible with the TSM mechanism appear as multiple nearby sequence changes and may be represented as combinations of SNPs, MNPs (single- and multi-nucleotide polymorphisms) and short indels in variant data. We showed earlier (Löoytynoja and Goldman 2017) that the TSM candidates identified between the two assemblies segregate in the 1000 Genomes population data; that TSM-like mutation patterns can be identified in genotype data; and that parent-offspring trios (Besenbacher et al. 2016) contain consistent de novo mutations (DNMs). Here, I revisit the topic using the haplotype-resolved data of Ebert et al. and the variation information compiled by massive sequencing studies (e.g., Karczewski et al. 2020; Taliun et al. 2021) and study the TSM mechanism’s role in generation of genomic variation. As haplotype-resolved genome data are still rare, I assess how reliably the TSM patterns found with denovo-assembled genomes can be identified using traditional short-read DNA sequencing.

## Results

### TSMs explain thousands of haplotypic mutation clusters

The SNVs, indels and SVs of the variant data of Ebert et al. (2021) (called here “HaplotypeSV data”; Table 1), were combined, and for each maternal and paternal genome copy, the clusters of sequence differences were identified (see Methods). For each mutation cluster locus within a haplotype, the identified variants were placed in their genomic background, creating two alleles with 150 bp of sequence context (Fig. 1A,B). Finally, having the two alleles – the region from the GRCh38 reference (called ‘REF’) and the alternative allele formed by the mutation cluster (called ‘ALT’) – the four-point alignment (FPA) algorithm (Löoytynoja and Goldman 2017) was applied to find the best solution involving a template switch, testing both alleles as the ancestral state (Fig. 1C,D; see Methods). Despite attempts to evaluate TSMs under a probabilistic model (Walker et al. 2021), it is unclear when a potential TSM solution should be considered more plausible than a combination of SNVs and indels. Here, the TSM candidates were required to: (i) be supported by at least two sequence changes of which at least one is a base change, and as a result show a higher sequence identity across the whole region than the original forward-aligned solution, and (ii) have a ②→③ region, inferred to be copied from the other template, of at least 8 bases long (Fig. S1). Despite strong evidence of many VNTRs (variable number of tandem repeats) evolving through template switching (Fig. S2), the candidate ②→③ regions were required to contain all four bases, thus removing hits within low-complexity sequences.

**Fig 1.**
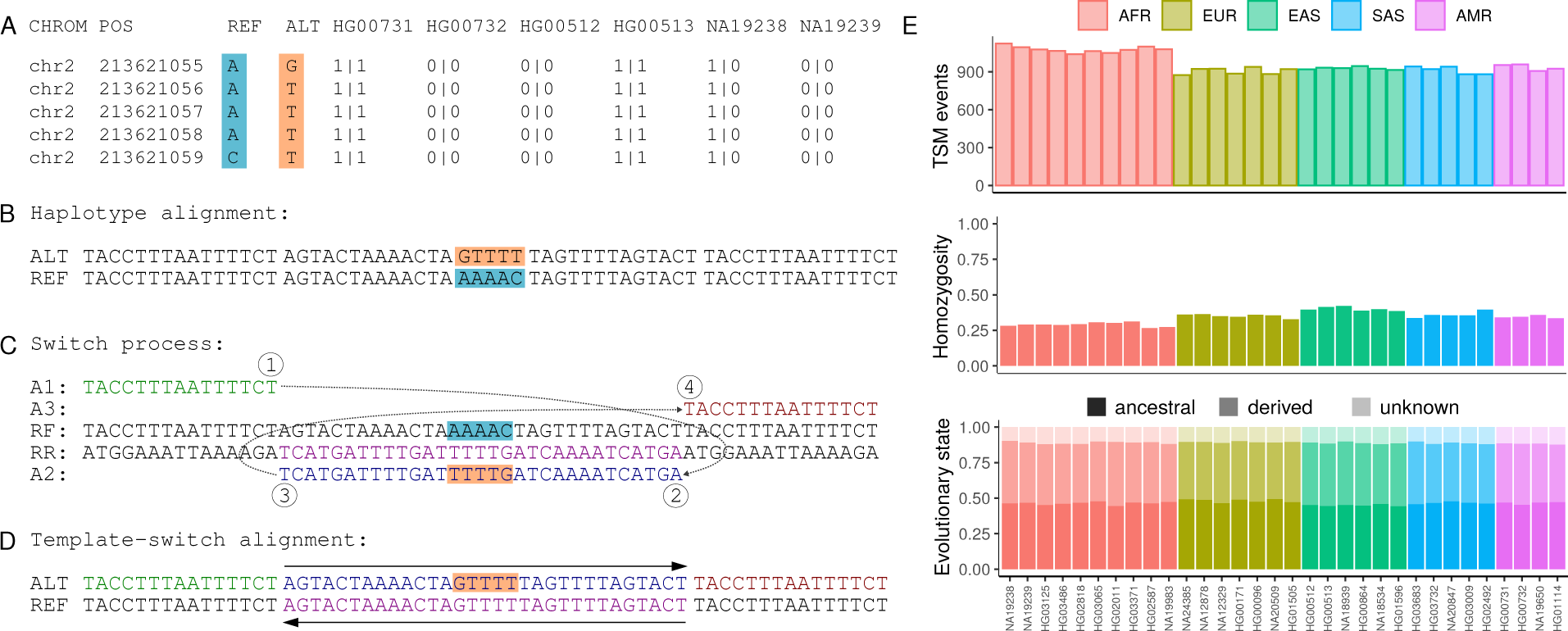
TSMs in HaplotypeSV data. (**A**) HaplotypeSV data contain a cluster of five SNVs that segregate as a unit among individuals (the first six are shown). (**B**) Within their genomic context, REF (cyan) and ALT (orange) alleles create two haplotypes. (**C**) The FPA algorithm attempts to explain the observed differences with two reciprocal template-switch events. In this case, the ALT sequence (A1, A2, A3; shown in green, blue and red) can be created from the REF sequence by copying the A2 fragment in reverse-complement (RR; shown in magenta). (**D**) The template-switch solution fully explains the cluster of five base differences. (**E**) Total numbers of inferred TSM events in different HaplotypeSV samples after removal of low-complexity sequences (top); proportion of homozygous loci (middle); and proportion of ancestral (dark), derived (intermediate) and unpolarized (light) alleles (bottom).

**Table 1.** Description of the datasets analyzed.

| Dataset | Source | Type | Platform | Read length | #samples | #variants |
| --- | --- | --- | --- | --- | --- | --- |
| HaplotypeSV | Ebert et al. | vcf | PacBio+Strand-seq | NA | 32 | 18,241,436 |
| Platinum NA12878 | Eberle et al. | vcf | Illumina | 100 PE | 1 | 4,167,900 |
| Chromium NA12878 | Weisenfeld et al. | vcf | Illumina Linked-Reads | 150 PE | 1 | 4,898,662 |
| 1kG phase 3 GRCh38 | 1000G Consortium | vcf | NA | NA | 2504 | 80,000,000+ |
| gnomAD.MNV full | Wang et al. | tsv | NA | NA | 125,000+ | 1,792,248 |
| gnomAD.MNV 3+ | Wang et al. | tsv | NA | NA | 125,000+ | 91,300 |
| second_gen.dnms | Sasani et al. | tsv | Illumina | 150 PE | 70 | 4,671 |
| third_gen.dnms | Sasani et al. | tsv | Illumina | 150 PE | 350 | 24,975 |
| Dataset | Source | Type | Platform | Read length | #samples | depth |
| Platinum NA12878 | Eberle et al. | bam | Illumina | 100 PE | 1 | 50x |
| Platinum CEPH 1463 | Eberle et al. | bam | Illumina | 100 PE | 17 | 50x |
| Chromium NA12878 | Weisenfeld et al. | bam | Illumina Linked-Reads | 151 PE | 1 | 56x |
| Chromium NA1289[12] | Marks et al. | bam | Illumina Linked-Reads | 151 PE | 2 | ~30x |
| PacBio GIAB NA12878 | PacBio (PRJNA540705) | bam | PacBio CCS Sequel II | NA | 1 | ~30x |

Per individual, 3,049–3,934 mutation patterns consistent with the TSM mechanism were identified, the longest of them 318 bp in length (Table 2, ‘all’). Many of these were in proximity of repeat elements and, due to mismapping of sequencing reads, could potentially be nonindependent events counted multiple times. After removing the loci masked as repetitive sequence in the reference genome, there were 872–1,121 high-quality TSM events per individual (Fig. 1E), the maximum length varying from 97-180 and the mean length being 11.56 bp (Table 2, ‘unmasked’; Fig. S3). Homozygosity of inferred TSM loci varied from 26.7–42.3% (Fig. 1E), and the proportion of singletons from 2.1–9.0% (Table S1). Although comparisons at the superpopulation level are affected by the different sample sizes, African (AFR) individuals expectedly had the lowest level of homozygosity and fixed loci, 29.2% and 0.7%, and the highest proportion of singleton loci, 7.8% (The 1000 Genomes Project Consortium et al. 2015), while those for Europeans (EUR, including Azkenazi NA24385) and Asians (AAS, including South and East Asians) were higher and lower, respectively (Table 2). The greater variation of African populations and the old age of the TSM patterns was reflected in the sharing of loci across the superpopulations: AFR shared 45.6% and 49.9% of its TSM loci with EUR and AAS, respectively, while the latter two shared 75.3–80.7% of their TSM loci with other superpopulations (Table 2). A unique feature of TSMs is that often alternative alleles can be polarized, i.e., have the ancestral and derived state determined, without an outgroup (Fig. S4). The ancestral allele could be defined in 87.7–90.3% of the cases and the proportion of loci with the ALT variant being the ancestral allele was found ranging from 49.7–55.3 across the individuals (Fig. 1E; Table S1). At the superpopulation level, Europeans had the highest average proportion of ancestral variants (Table 2), probably reflecting the European origin of the GRCh38 reference genome and thus capturing a greater amount of derived European TSM variation. Considering the candidate TSMs in the unmasked part of the reference genome, there were in total 2,200 unique loci. While the annotation revealed 96.2% of the mutation loci to be intergenic or intronic (Table S2), 24 loci were inferred as coding and I had a closer look on three cases that had the ②→③ region of at least 12 bp long. Segregation of the ALT variants revealed one of these as a false positive, but the two others seemed real. The inferred TSM events created a cluster of linked base substitutions and indels in the variant data and appeared at the lowest possible frequency, heterozygous in a single individual; the mutations resulted in an early stop codon in an alternatively-spliced exon of gene *ANP32E* (Fig. S5) and changes in a nonsense-mediated decay destined transcript of gene *PHF21A*.

**Table 2.** TSM statistics for HaplotypeSV data.

| Superpopulation<br>Haplotypes |  | AFR<br>20 | EUR<br>14 | AAS<br>22 |
| --- | --- | --- | --- | --- |
| ALL | TSMs/individual | 3783 | 3135 | 3173 |
|  | unique TSMs | 12,501 | 7,370 | 8,956 |
|  | maximum length | 318 | 188 | 188 |
|  | average length | 13.14 | 13.14 | 13.15 |
| UNMASKED | TSMs/individual | 1,074 | 906 | 919 |
|  | unique TSMs | 3,133 | 1,899 | 2,246 |
|  | maximum length | 180 | 180 | 180 |
|  | average length | 10.61 | 10.56 | 10.55 |
|  | singletons | 7.8% | 2.7% | 3.2% |
|  | fixed | 0.7% | 1.8% | 2.8% |
|  | homozygous | 29.2% | 35.3% | 38.5% |
|  | ancestral | 52.0% | 53.9% | 51.5% |
|  | shared with AFR | — | 75.3% | 69.6% |
|  | shared with EUR | 45.6% | — | 68.2% |
|  | shared with AAS | 49.9% | 80.7% | — |

### Known TSM loci can be genotyped with short-read data

One individual in the HaplotypeSV data, NA12878, is the mother of a three-generation 17-member pedigree (CEPH 1463) sequenced to high coverage (Eberle et al. 2017) as well as one of the reference samples of the Genome in a Bottle (GIAB) initiative (Zook et al. 2016). The TSM loci identified in NA12878 were genotyped in different GIAB datasets and in the pedigree 1463 to assess their de novo vs. standing, in vitro vs. in vivo, and somatic vs. germline origin. Using HaplotypeSV genotype data for NA12878, the genome contexts for the alternative alleles were reconstructed and, for each individual in the family, the reads extracted from that particular loci were mapped against the two alternative alleles. Outside the mutation locus, short reads should map evenly between the two alleles, but reads that overlap the positions differing between the alleles are expected to map to one allele only (Fig. 2A,B). By comparing the read coverage across the two alternative alleles, the test individual could be identified as homozygote for REF allele (R*|*R), heterozygote (R/A), or homozygote for ALT allele (A*|*A). A strategy similar to base calling was adopted and a 99% posterior probability was required for credible genotype calls (see Methods; Fig. S6).

**Fig 2.**
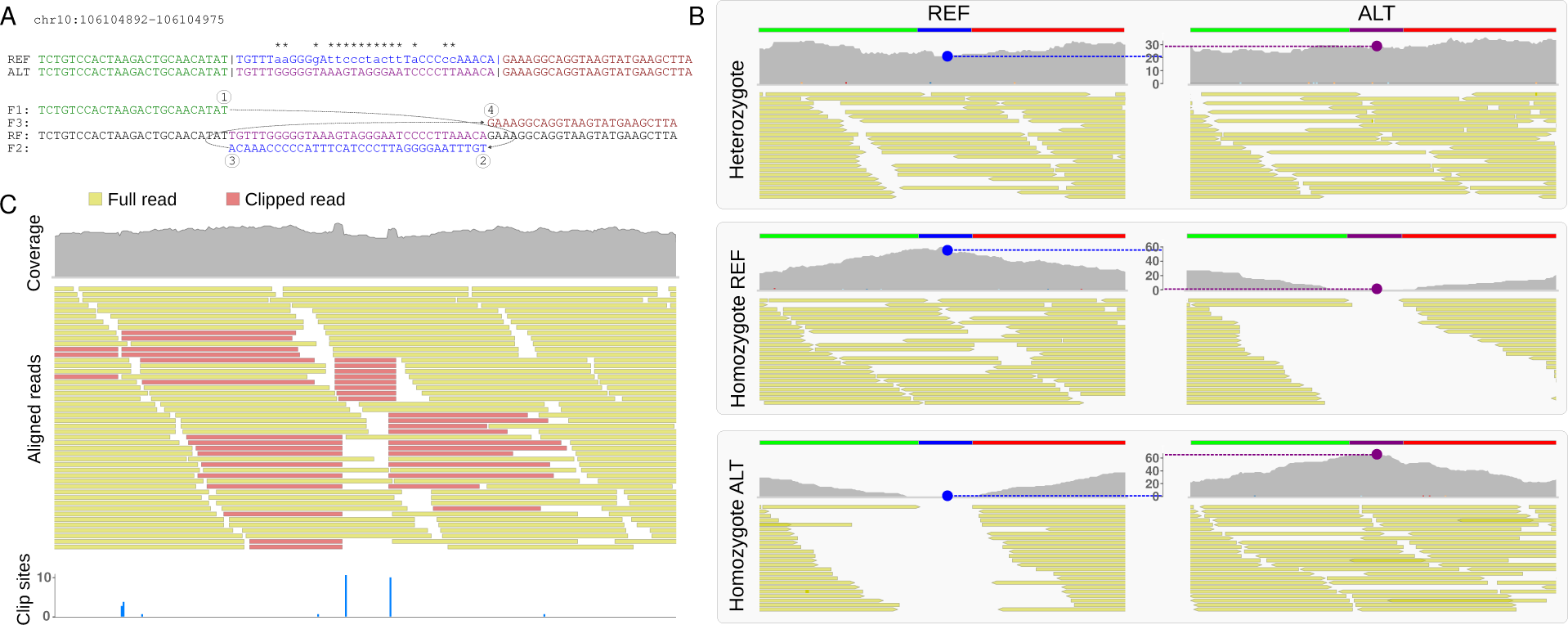
Genotyping of a TSM locus. (**A**) TSM solution for a complex mutation in chr10. The REF and ALT alleles share the left (green) and right (red) flanking region but have a different central part (blue and magenta). The differences in the central part are explained by a TSM event (below). (**B**) The two haplotypes with alternative central parts (blue and magenta bars) are used as the reference for re-mapping of reads extracted from the locus. In a heterozygous individual (top), reads are mapped evenly to the two alleles, giving uniform coverages (gray). In individuals homozygous for the REF allele (middle) and ALT allele (bottom), reads covering the central part are mapped predominantly to one allele only. Genotype is inferred of the mapping coverage in the middle of the locus (blue and magenta dots and dotted lines) using a function similar to that for genotype calling on nucleotides. (**C**) NA12878 Platinum data show fairly normal mapping coverage (gray, top) but a closer look on the reads (full and clipped reads in yellow and red, respectively) reveals an anomaly. The cluster of clip sites (blue; bottom) allows computational identification of the locus.

I started with the Illumina Platinum (Eberle et al. 2017) and 10x Genomics Chromium (Weisenfeld et al. 2017; Marks et al. 2019) datasets for NA12878 (Table 1) and genotyped the TSM-like loci found in the HaplotypeSV data. Focusing on the 864 loci that were not masked as repeats or low-complexity sequences and had enough read coverage in both data sets, the two short-read datasets were found to provide the genotype and agree on it in 831 cases (Fig. 3A; Table S3). While 14 loci could not be genotyped with the short-read data using the 99% posterior probability cutoff, the variant pattern in the HaplotypeSV data was inconsistent in 84 cases, and the genotype of the loci could not be determined. Of the 780 loci inferred to be either heterozygous or homozygous for the ALT allele in the HaplotypeSV data, the three datasets agreed in 735 cases (Table S3). Eight loci recorded variable in the HaplotypeSV data were inferred to be homozygous for the REF allele in both short-read datasets. Many of these were in a repetitive sequence context and all loci were found to segregate among other samples in the HaplotypeSV data, ruling out the possibility of DNMs. Although the reasons for inconsistent HaplotypeSV genotypes could not be resolved, 74 of 84 cases showed intermediate depth ratios in short-read data, i.e. *r* was not *∼* 1, *«* 1 or *»* 1 (Fig. 3A; orange symbols around -0.5, -0.5). If these were de novo mutations, one would expect the ratios to be reverse, and a more likely explanation is cross-mapping from duplicated sequence blocks. A small number of candidate TSM loci showed inconsistency between different datasets, possibly due to similarly incorrectly mapped reads (Fig. S7).

**Fig 3.**
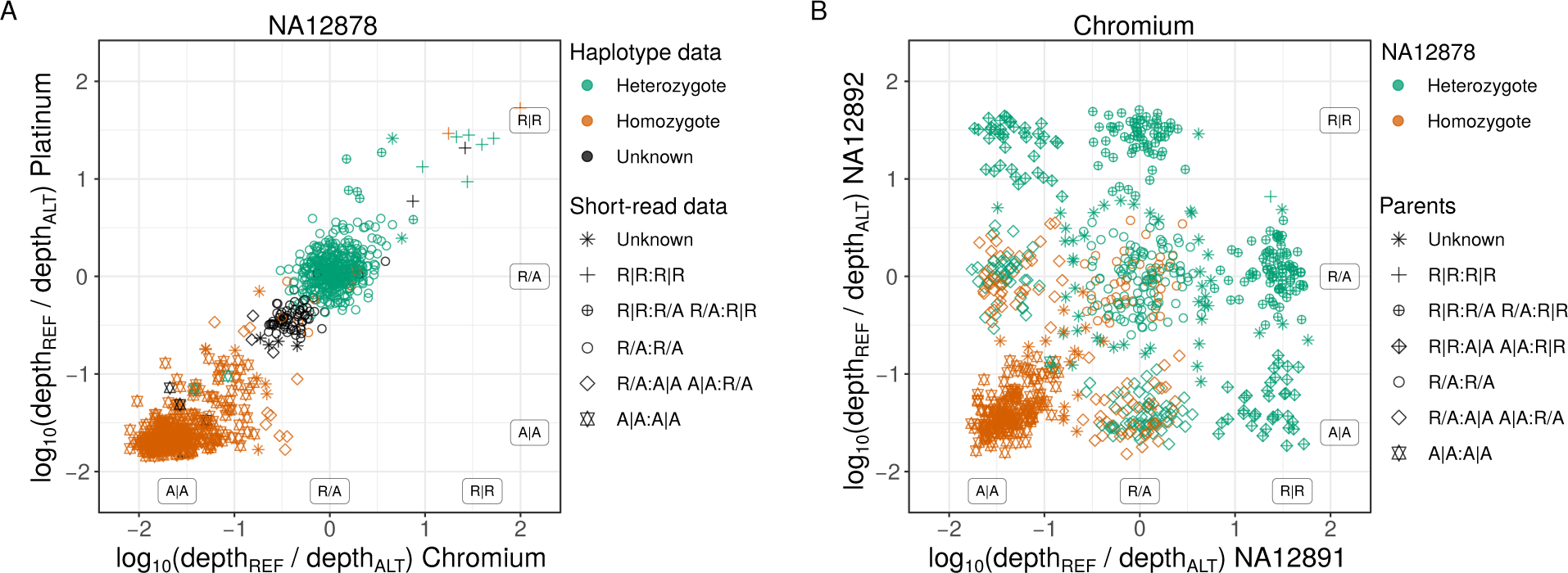
Confirmation of HaplotypeSV TSM loci with short-read data. (**A**) Ratio of REF and ALT allele mapping coverage (*r* = *depth*_REF_*/depth*_ALT_) reflects the genotype: *r ≈* 1 (and thus *log*_10_(*r*) *≈* 0) for a heterozygote; *r «* 1 and *r »* 1 for the two types of homozygotes. The Log10-ratios agree for the NA12878 Platinum and Chromium datasets, and 95.5% of the 883 loci identified in HaplotypeSV data are called to contain at least one ALT allele with 99% posterior probability in both datasets; 17 and 10 loci are called homozygous REF with at least one dataset and by both (top-right corner). (**B**) Of the 783 variable loci in NA12878 Chromium data, all but one locus are called to contain at least one ALT allele in the parents. In the 290 loci homozygous for ALT in NA12878 (orange), both parents contain at least one ALT allele. ’Unknown’ indicates inconsistent variant data or posterior probability below 0.99. One pseudo-count was added to all values to avoid divisions by zero. In legend, the inferred genotypes for the two datasets are separated by a colon, and X_1_*/*X_2_ represents the alternative arrangements of the two alleles, X_1_|X_2_ and X_2_|X_1_.

**Table 3.** Genotypes (%) of TSM loci in CEPH 1463 children.

| Parental<br>genotypes | n | Offspring expected |  |  | Offspring observed |  |  |  |
| --- | --- | --- | --- | --- | --- | --- | --- | --- |
|  |  | R\|R | R/A | A\|A | R\|R | R/A | A\|A | NA * |
| R\|R,R\|R | 7 | 100 | 0 | 0 | 72.7 | 24.7 | 0 | 2.6 |
| R\|R,R/A | 169 | 50 | 50 | 0 | 48.6 | 50.5 | 0.5 | 0.4 |
| R\|R,A\|A | 37 | 0 | 100 | 0 | 0 | 98.5 | 1.5 | 0 |
| R/A,R/A | 214 | 25 | 50 | 25 | 16.8 | 63.5 | 17.5 | 2.2 |
| R/A,A\|A | 206 | 0 | 50 | 50 | 0.4 | 53.0 | 45.1 | 1.6 |
| A\|A,A\|A | 177 | 0 | 0 | 100 | 0 | 0.5 | 98.5 | 1.1 |

\*Loci with posterior probability below 0.99 considered as unknown

To further verify the genotyping approach, I performed a similar analysis on the parents and children of NA12878. On the parents, I genotyped the 783 loci found in the HaplotypeSV data and called to contain at least one ALT allele in the NA12878 Chromium data. The inferred genotypes showed strong agreement and, of the 307 loci called homozygous for the ALT allele in NA12878, 290 could be genotyped with Chromium data of both parents and each of them contained at least one ALT allele (Fig. 3B; Table S4). The only locus that appeared as a novel mutation according to parental data (R*|*R:R*|*R in Fig. 3B) was a false positive and in manual verification, one parent was found to contain two supporting reads. Genotyping of the loci in the 11 children (CEPH pedigree 1463) required their transfer from GRCh38 to GRCh37. The lift-over worked and the parents’ (NA12877 and NA12878) alignment data (Platinum/GRCh37) contained enough reads to genotype 843 of the TSM-like loci found variable in NA12878 in the HaplotypeSV data. Except for loci that were inferred to be heterozygous in both parents (R/A,R/A), the observed genotypes matched nearly perfectly the expectation under the Mendelian segregation ratio (Table 3). An excess of heterozygotes in genotype data is a classic mark of artifact caused by sequencing reads originating from different loci, e.g., due to duplication of genome regions. Seeing those among the CEPH 1463 children suggests that a proportion of inferred variants for NA12878 in the HaplotypeSV data are erroneous: the TSM events causing those variants may be real, but they have happened in duplicated copies, not within the loci where the calls were made.

### Novel TSM loci can be discovered with short-read data

Many of the TSM loci detected in the HaplotypeSV data had not originally been called with the short-read data, despite the two alleles differing over several tens of bases. A closer look at these revealed that alternative TSM haplotypes can show nearly uniform sequencing coverage across the region if the ②→③ region of the inferred TSM process (Fig. 2A) is long enough: the mapping algorithm then produces a decent mapping for the read core by clipping the mismatching overhangs (Fig. 2C). Although this is clearly undesirable behavior, the observation suggested a strategy to identify potential TSM loci in short-read data by searching for clusters of clip sites, extracting the reads mapped to the region, and then producing a local reassembly of the reads; in positive cases, the dissimilarities between the assembled sequences could be explained by a TSM event.

To test that, I took SvABA, a tool developed for the detection of structural variants by local reassembly of short sequencing reads (Wala et al. 2018), and integrated the FPA algorithm (Löoytynoja and Goldman 2017) to search for TSM patterns in the resulting contigs. Clusters of clip sites were identified in short-read data and clusters of variants in the corresponding variant data, performing the analysis independently on the Illumina Platinum (Eberle et al. 2017) and 10x Genomics Chromium (Weisenfeld et al. 2017) resources for NA12878. SvABA-FPA was then applied on each candidate locus (see Methods), and 1,054 and 26,222 candidate TSM loci were found in the Platinum and Chromium data, respectively. Of these, 211 and 755 loci, respectively, passed the sequence-based filtering and 203 and 299 loci the subsequent mapping depth-based genotyping (being either heterozygous or homozygous for the ALT allele, see Fig. S8 and Methods). The fact that 456 of the 755 Chromium hits passing the first stage of filtering could not be confirmed with read mapping indicates that the alternative haplotypes were created by a small number of reads. Whether these reads had originally been misaligned or represent low-frequency TSM mutations, possibly originating in vitro, could not be confirmed with current data; similar hits not found in the Platinum data may be explained by different read lengths (Table 1), technological differences between the standard and linked-reads Illumina sequencing and downstream bioinformatic methods used (Li 2013; Marks et al. 2019). Reassuringly, a majority of the loci passing all filtering, 89.7% and 70.6% for Platinum and Chromium, respectively, were shared either with the other short-read approach or with the HaplotypeSV set (Fig. 4). On the other hand, the large number of candidate loci, 690 of the total 843, identified uniquely with the HaplotypeSV data suggests that a great majority of the loci do not contain misaligned and soft-clipped reads: if no variants were called at those loci in the Platinum and Chromium sets, no signal was present to include the loci for the short-read based TSM search.

**Fig 4.**
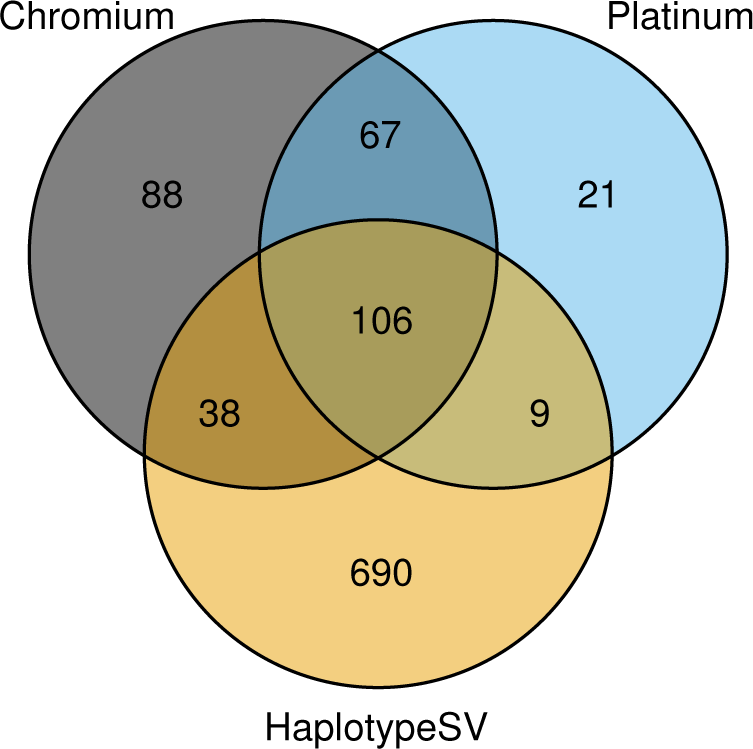
The number of TSM candidate loci identified in different data. The diagram shows the overlap of TSM candidate loci identified using the Platinum and Chromium short-read data and the HaplotypeSV genotype data for the reference individual NA12878. The Platinum and Chromium loci have passed sequence-based filtering and all sets have passed the mapping depth-based genotyping.

I studied in more detail 59 cases that had the ②→③ fragment of at least 25 bases long (Data S1). Most of the cases showed the expected signal in the original alignment data that was removed when the same reads were mapped against the two alternative haplotypes (Fig. 5A,B) while in a few rare cases the variant calling had correctly captured the differences between the two TSM haplotypes (Fig. 5C,D). Some of the heterozygous loci were not called in any variant call set despite significant differences between the two haplotypes (Fig. S9, Data S1), and, unexpectedly, several loci homozygous for the ALT allele were uncalled in different variant call sets, including the HaplotypeSV data (Fig. S10). As an example of the latter, 67 TSM candidates found in, and confirmed by (Fig. S8D), both Platinum and Chromium data were not present in the HaplotypeSV set. I had a closer look on the 16 loci with the ②→③ fragment of at least 25 bases long, and examined the loci in PacBio long-read sequencing data and 1000 Genomes variantion data (The 1000 Genomes Project Consortium et al. 2015; Table 1; Data S2). All the studied cases were confirmed to be genuine mutation clusters by the PacBio data, and two types of errors in HaplotypeSV calls were observed: the representation of the mutation cluster was incomplete, thus not allowing correct reconstruction of the the alternative allele and discovery of the TSM event (Fig. S11A); and complete lack of variant calls (Fig. S11B). In an extreme, the variants were missing from HaplotypeSV data despite being called in full for NA12878 in the 1000 Genomes data and clearly supported by all sequencing data (Fig. S11C).

**Fig 5.**
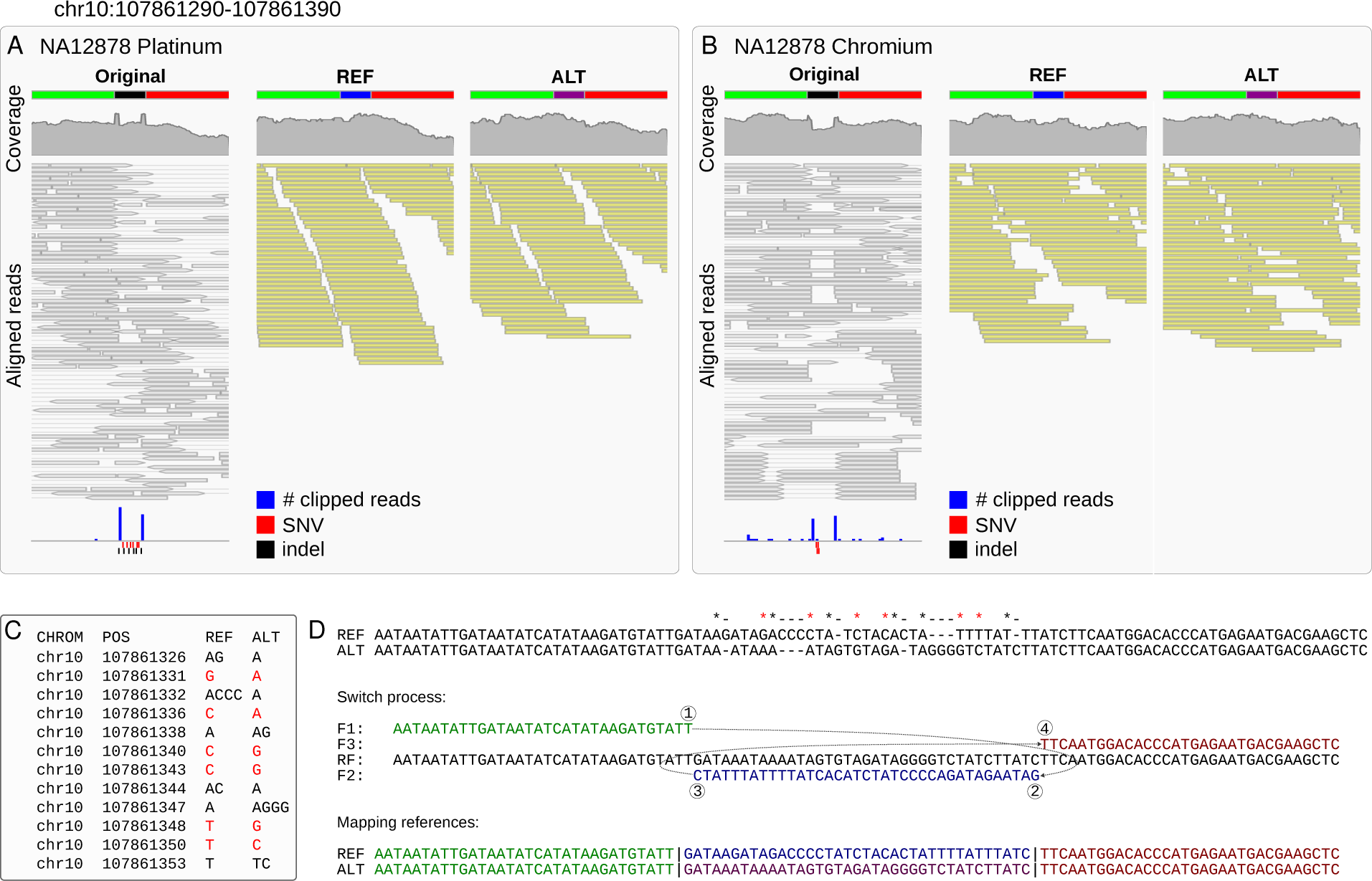
TSM candidate identified with short-read data. (**A**) In the Platinum data (left), a region in chr10 has excess of soft clips (bottom, blue bars) and called variants (SNVs in red, indels in black); the mapping coverage (top, in gray) also shows atypical patterns. (**B**) Similar signals are seen in Chromium data. (**C**) The Platinum variant data contain six SNVs and six indels within 28 bp. (**D**) De novo assembly of the reads creates two locally highly dissimilar haplotypes compatible with the called variants (top). All differences can be explained with a TSM event, an inversion in place (middle). Using two haplotypes with alternative central parts (blue and magenta; bottom) as the reference, extracted reads map in full length (**A,B**; middle, right) with roughly even coverages. No variants for NA12878 were called in this region in HaplotypeSV data.

### TSMs explain thousands of clusters in variation databases

A significant fraction of the TSM candidate loci were missing from the original variant call sets (Fig. 5C) – 16 of the 59 closely studied loci were missing from both sets (Data S1) – but none of the manually confirmed cases supported by both short-read datasets was completely absent from the dbSNP database (Sherry et al. 2001). Although dbSNP correctly lists many studied loci as multinucleotide variants (MNV), all loci were also present as multiple SNVs and indels and some only as multiple independent MNV or indels (Fig. S12, Data S1). The latter is slightly surprising as variants originating from a complex mutation are expected to be present in their entirety or not at all, and similar allele frequencies should allow phasing and grouping independent variants into MNVs (Choi et al. 2018).

The Genome Aggregation Database (gnomAD, v2.1.1; Karczewski et al. 2020) contains variation found among 125,000 exomes and 15,700 genomes of unrelated individuals sequenced as part of various disease-specific and population genetic studies. Wang et al. (2020) identified gnomAD SNVs appearing within proximity of 1-10 bp, and provide them as phased variant pairs (Table 1). Using these data, I selected variant pairs with similar allele counts (ACs; ΔAC *≤*10%), merged the data of different variant-pair distances, and for all clusters of at least three variants, created the alternative alleles, and searched for a TSM solution to explain the differences (see Methods). Among the 91,300 clusters of three or more SNVs, I was able to find a TSM solution for 4,425 loci. Among these, I had a closer look at the 192 cases where the ②→③ region was at least 25 bp long and the REF and ALT alleles differed at least by four SNVs (Data S3).

Among the studied cases, the TSM mechanism perfectly explained MNVs consisting of up to 15 base changes (Fig. 6) while the longest inferred TSM events had the ②→③ region of over a hundred bases (Fig. S13). Consistent with previous findings, 36% of the TSM loci studied contained low complexity sequence or VNTRs, but base changes, including those in the two longest events of 119 and 125 bp (Fig. S13B,C), were not necessarily explainable by a simple slippage mechanism; within a complex sequence, the longest TSM event was 90 bp long (Fig. S13A). A locus in chromosome 19, present in a single individual, was of particular interest as 16 of the 19 base changes of gnomAD MNV could be explained with two adjacent 62 and 32 bp TSM events separated by a 16 bp forward fragment (Fig. S14). This indicates that, similarly to larger arrangements (Lee et al. 2007; Zhang et al. 2009), local TSMs can also be chained into complex combinations. One should note that the gnomAD MNV data lack indels and no TSM patterns involving sequence length changes were included, meaning that the real number of mutation clusters explainable by the TSM mechanism is likely much higher.

**Fig 6.**
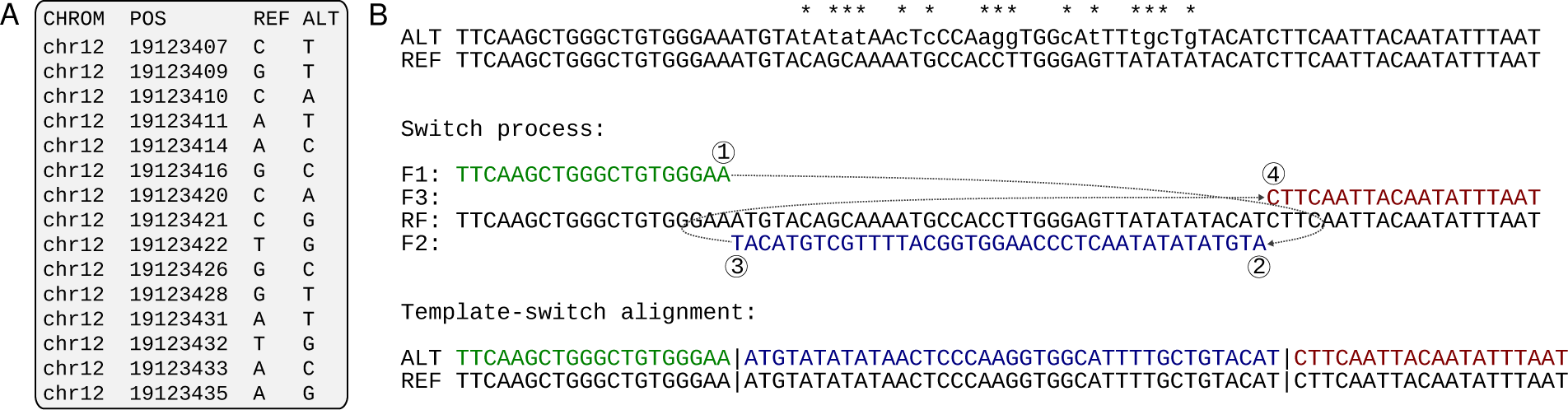
Complex gnomAD MNV explained by a single TSM. (**A**) Wang et al. (2020) identified 15 SNVs within a 29 bp interval in chr12. (**B**) All the differences (top, marked with asterisks) can be explained with a TSM inverting 39 bp-long fragment in place (middle, bottom).

I replicated the analysis with the DNM data from 33 large, three-generation CEPH families by Sasani et al. (2019; Table 1). Although no TSM-like patterns were found among the germline DNMs of the 70 second-generation individuals, four consistent patterns (Fig. S15) were observed among the 24,975 de novo SNVs and small indels of the 350 third-generation individuals. As most variants were isolated and there were only 218 DNM clusters, this gives a frequency of 1.8% for TSMs among the clustered DNMs. Sasani et al. estimated that, on average, there are 70.1 de novo SNVs and 5.9 de novo indels per genome, but the germline status of third-generation variants cannot be verified. In general, the vast majority of mutations are somatic, Conrad et al. (2011) estimating a ratio of 20:1 for non-germline to germline DNMs, and the observed TSM-like patterns are also likely to be somatic.

## Discussion

My analyses extend the previous works (Löytynoja and Goldman 2017; Walker et al. 2021) and show that human genomes have thousands of mutation patterns consistent with DNA replication-related template switching and that these loci segregate within and between populations. I studied the haplotype-resolved genotype data by Ebert et al. (2021), two independent WGS experiments on the reference individual NA12878 and her extended family (Eberle et al. 2017; Weisenfeld et al. 2017; Marks et al. 2019), and variant information in the gnomAD database (Karczewski et al. 2020; Wang et al. 2020) and from a DNM study (Sasani et al. 2019). While I found Ebert et al.’s HaplotypeSV data to have captured large numbers of TSM patterns and mostly reflect the haplotypic differences between the chromosomes, the call set was far from perfect and I could identify many additional TSM-like loci using the independent short-read datasets. The characteristics of the loci missing from the HaplotypeSV data raise serious questions about the completeness of the Ebert et al.’s variant set, and some of the omitted TSM-like mutation patterns were perfectly visible in the sequencing data and called in full for exactly the same individual in the 1000G variant data. In addition to short-read data, I identified de novo TSMs in data of Sasani et al., though was not able to confirm their germline status, and saw consistent singleton TSM patterns in gnomAD variant data (Karczewski et al. 2020; Wang et al. 2020). Overall, my results demonstrate that, despite their relatively low rate, the TSM mechanism explains a significant fraction of MNVs seen in human variation data and thus contributes to correlation in local mutation frequencies (Śegurel et al. 2014; Harris and Nielsen 2014).

While the ability of TSMs to create reverse-complement copies of short sequence fragments is highly interesting, the mechanistic origin of sequence differences is irrelevant to many applications of genome analysis. My analyses revealed that TSM-like mutations are found within genes and other potentially functional elements, and there are anecdotal reports of TSM-like mutations causing genetic diseases in humans (Menardi et al. 1997). Given this, a worrying finding of the study was that the DNA sequencing pipelines capable of identifying SNVs may miss inverted fragments of a few tens of bases. The difficulty of detecting certain TSM patterns comes from the ability of mapping tools to align the affected reads in their expected context after excessive clipping of the read ends. If this mapping error goes unnoticed in variant calling, it can invalidate e.g. medical genetic studies searching for causative alleles behind genetic diseases. On the other hand, Seplyarskiy et al. (2021) recently proposed an approach to separate the signals of different mutation processes, and among others, described signals from “asymmetric resolution of bulky DNA damage” and from “asymmetric replication errors”. Such asymmetric signal could be created by template switching if the 10 times higher frequency of TSMs in leading strand seen bacteria (Rosche et al. 1997; Seier et al. 2011) applies to other organisms, or if the mechanism is related e.g. to coordination or clashes of transcription and replication systems (Hamperl et al. 2017; Chen et al. 2019). Local TSMs have many similarities with the FoSTeS (Lee et al. 2007) and MBIR (Hastings et al. 2009) mechanisms – and possibly with chromoanasynthesis in cancer (Liu et al. 2011; Holland and Cleveland 2012) – but due to their short length they are expected to be more benign (Zhang et al. 2009). With my novel tools TSM patterns can be identified in different types of genome data, enabling analyses of their genome-wide distribution and possible correlation with different cellular processes (Seplyarskiy et al. 2021). Such analyses should bring us closer to understanding the mechanisms underlying the template switching.

## Methods

### Data

Ebert et al. genotype data (VCF; freeze3) were downloaded from http://ftp.1000genomes.ebi.ac.uk/vol1/ftp/datacollections/HGSVC2/release/v1.0/integrated callset, combining SNV, indel and SV calls. Chromium data (BAM;VCF) were downloaded from http://ftp-trace.ncbi.nlm.nih.gov/giab/ftp/data/NA12878/ 10Xgenomics ChromiumGenome LongRanger2.0 06202016 and http://ftp-trace.ncbi.nlm.nih.gov/giab/ftp/technical 10Xgenomics ChromiumGenome LongRanger2.0 06202016. Platinum data (BAM;VCF) were downloaded from http://ftp.1000genomes.ebi.ac.uk/vol1/ftp/datacollections/illumina platinum pedigree/data/CEU, via http://github.com/Illumina/PlatinumGenomes, and from dbGaP (http://www.ncbi.nlm.nih.gov/projects/gap/ cgi-bin/study.cgi?study id=phs001224.v1.p1; authorized access for A.L.). Pacific Biosciences contribution to NIST GIAB initiative was downloaded from https://ftp-trace.ncbi.nlm.nih.gov/ReferenceSamples/giab/ data/NA12878/PacBio SequelII CCS 11kb. 1000 Genomes variant data were downloaded from http://ftp. 1000genomes.ebi.ac.uk/vol1/ftp/datacollections/1000G 2504 high coverage/working/phase3 liftover nygc dir/. gnomAD data were downloaded via https://gnomad.broadinstitute.org/downloads#v2-multi-nucleotide-variants and Sasani et al. data through https://github.com/elifesciences-publications/ceph-dnm-manuscript/tree/ master/data. Reference genomes were downloaded from http://ftp.1000genomes.ebi.ac.uk/vol1/ftp/data collections/HGSVC2/technical/reference (GRCh38) and from http://ftp.1000genomes.ebi.ac.uk/vol1/ftp/ technical/reference (GRCh37 and 38). Description of the datasets is given in Table 1. With the exception of the CEPH 1463 children, all data used in this study are publicly available and the instructions for their download and analysis are provided in the Supplementary Code.

### Analysis of haplotype data

The genotype data of Ebert et al. (i.e., HaplotypeSV data) were processed with a custom Python script (tsm scan SV2.py) internally utilising BWA (Li 2013), SAMtools and BCFtools (Danecek et al. 2021). The script detects clusters of variants and creates alternative haplotypes by replacing parts of the reference genome sequence with the variant bases. The reconstructed haplotypes (with 150 bp of flanking sequence) are then compared to the reference allele, and a TSM solution is searched reciprocally to create one of the alleles from the other. Each individual and chromosome was processed independently. The candidate loci in NA12878 were processed with another Python script (tsm alleles2.py) that (i) creates the two alleles for each locus using the reference sequence and the HaplotypeSV variant information, (ii) extracts the reads mapped to the locus in a BAM alignment file and maps these reads to the two alternative alleles, and (iii) computes the mapping coverage statistics for the two alleles, recording the mapping depth at the region differing between the alleles as well as immediately upstream and downstream of the differing region. The REF allele was taken from the reference genome, and the ALT allele was created by placing the NA12878 variants (each haplotype separately) into a copy of that; the differing regions were then placed in identical context with 500 bp of flanking sequence from the reference genome.

The genotype of the locus was inferred from the mapping coverage of the two alternative alleles using a strategy similar to base calling. I required a minimum coverage of 10 reads for upstream, downstream and within the differing region for either of the alleles. Then the probabilities for the three possible genotypes were obtained as:

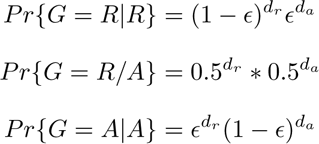

where *e* is the error rate of 0.01 for the read being mapped to a wrong allele, and *d_r_* and *d_a_*are the mapping coverages for the REF and the ALT alleles. The inferred genotype *X*, where *X ∈ {R|R, R/A, A|A}* (standing for homozygote REF, heterozygote and homozygote ALT, respectively), was then the genotype with the highest probability if that was at least 99% of the total probability:

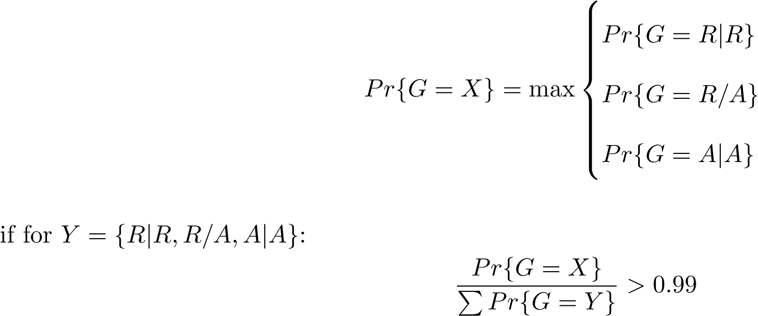

The variant annotation was performed using the R package VariantAnnotation (Obenchain et al. 2014) and the software BEDTools (Quinlan and Hall 2010) based on Ensembl v. 106 gene models (Homo sapiens. GRCh38.106.chr.gff3). Only one count of each annotation class was considered for each locus.

### Analysis of short-read data

The FPA algorithm (Löoytynoja and Goldman 2017), written in C++ and available through https://github.com/ariloytynoja/fpa, was integrated into the SvABA tool (Wala et al. 2018). Clusters of more than one base mismatch were identified in VCF data using BCFtools and an awk script counting variants within 20-bp intervals. Similarly, clusters of more than ten clipped reads were identified in BAM data using SAMtools and an awk script calculating the positions of soft clips based on the CIGAR string and another script identifying clusters of positions within 20-bp intervals. Candidate loci were targeted with SvABA-FPA and, for contigs showing two or more base differences in comparison to the reference sequence, TSM solutions were computed using both the REF and the ALT allele (i.e. SvABA-created contig) as the ancestral type. The resulting TSM candidates were filtered and those overlapping with repeat elements or low-complexity sequence, or having short length or low identity at flanking sequences were removed. More precisely, loci intersecting with masked sequence or assembled contigs having low sequence complexity (Trifonov’s complexity with order 5 *>* 0.25; computed with program SeqComplex (Caballero et al. 2014)) were removed; TSMs were compared to forward alignments and required to be better, containing at least two edit events (of which at least one base mismatch) less than the non-template-switching alignment; of those, ones with ① − ④ distance longer than 250 bp or shorter than 5 bp and upstream/downtream flanking regions or ②→③ fragment showing sequence identity below 95% or being less than 10 bp long were discarded.

For the loci passing the sequence-based filtering, the REF and ALT alleles were placed in identical context with 500 bp of flanking sequence. Alignment data (in BAM format) were genotyped by extracting the reads and mapping them to the two alleles using the custom Python script. The genotype was inferred from the read coverage as explained above, and loci inferred as heterozygous or homozygous for the ALT allele were retained. Overlapping loci were counted using the R package GenomicRanges (Lawrence et al. 2013). The custom scripts for all steps and instructions for their use are provided in the Supplementary Code.

### Analysis of CEPH 1463 data

The coordinates of candidate loci were transferred from GRCh38 to GRCh37 with CrossMap v0.2.9 (Zhao et al. 2014) using the Ensembl chain file. The same methods were used to genotype the loci.

### Analysis of MNV data

The MNV data from gnomAD and by Sasani et al. were processed similarly to the HaplotypeSV data with a Python script (tsm scan dnm.py). The script detects clusters of variants and creates alternative haplotypes by replacing parts of the reference genome sequence with the variant bases. It then compares the reconstructed haplotypes (with 150 bp of flanking sequence) with the reference allele and, reciprocally, searches for a TSM solution to create one of the alleles from the other. Tandem repeats were searched with trf (Benson 1999) and hits containing them in ②→③ region were discarded.

### Software availability

The Python/awk scripts used, the source code for the SvABA-FPA tool, and the instructions for their use are provided as Supplemental Code and deposited on GitHub at https://github.com/ariloytynoja/short-range-template-switching.

### Ethics statement

A.L. had authorized access to the dbGaP genome data for the CEPH 1463 offspring (http://www.ncbi.nlm.nih.gov/projects/gap/cgi-bin/study.cgi?study id=phs001224.v1.p1) under the project name “Properties of de novo template switch mutations”. The offspring data were analysed in a secure computer environment accessible only by the author, and only summary statistics of genotype inheritance are reported.

## Acknowledgments

I thank CSC – IT Center for Science and the University of Helsinki IT Center for the computing resources and the secure analysis environment, and the UH Biodata Analytics Unit for helpful discussions. This work was enabled by the Academy of Finland grant #322681.

---

**Fig S1.**
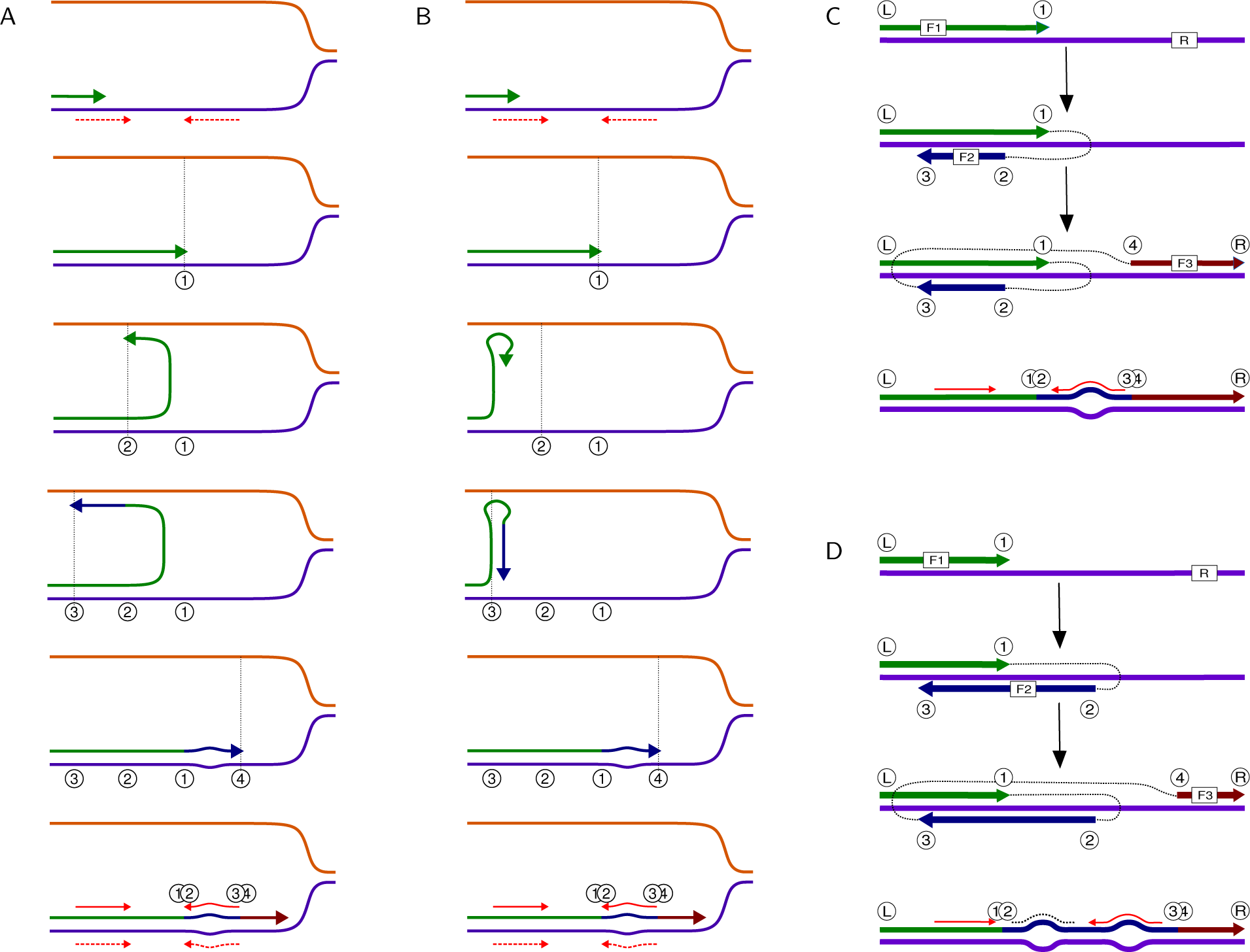
Classic vs. four-point model of template switching. (**A**, **B**) In the classic model of Ripley (1982), the replication (green arrow) switches the template and starts to go backward either along the opposite strand (**A**) or along the newly synthesized strand (**B**). After a synthesis of a short fragment (blue arrow), the replication switches back to the original strand and continues normally (dark red arrow). As a result, the near-perfect inverted repeat (red dashed arrows) has been converted into a perfect inverted repeat (red arrows), causing mismatches between the sequences (bulge). (**C**) The four-point model of Loöytynoja and Goldman (2017) is a computationally tractable representation of the switch process and relaxes some of the assumptions underlying the Ripley model. The model assumes four points, ① − ④, where the replication process changes from one template to another and copying of the fragment ②→③ in a reverse complement manner. The outcome of the switch process depends on the relative positions of points ① − ④, and the type 3-2-1-4 (read from left to right) shown here is consistent with the process shown in (A) & (B) and can take place either intra- or inter-strand. (**D**) The outcome of the template switch process depends on the relative positions of points ① − ④, and certain combinations, such as 3-1-2-4 shown here, can occur only interstrand, some may produce multiple repeats, and others only an inversion. See Löoytynoja and Goldman (2017) for enumeration and outcomes of all possible switch point combinations. In this study, candidate template switch events were required to explain at least two sequence changes (indicated with bulges in the schematic figures) of which at least one is a base change, and have ②→③ region, inferred to be copied from the other template, of at least 8 bases long. It is possible that some of inferred template switch events were in fact created by multiple independent base changes, insertions or deletions.

**Fig S2.**
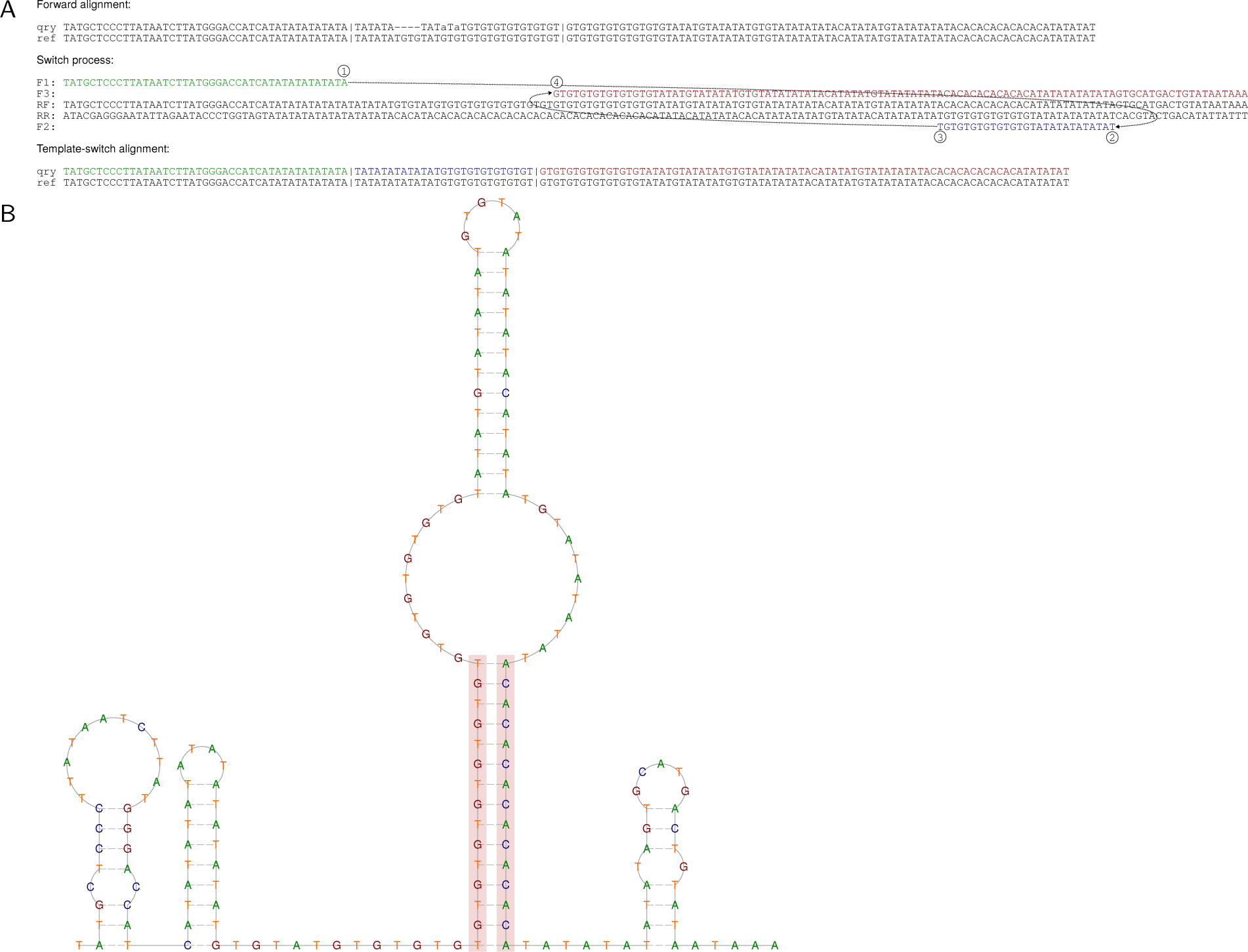
Changes in a VNTR locus consistent with the TMS process. (**A**) The differences between the alternative allele (qry) and the GRCh38 copy (ref; top) at chr1:116324126-116324132 can be explained by template switching (middle, bottom). The same outcome could be obtained with two slippage mutations, however. (**B**) DNA secondary structure reveals that, in addition to runs of internally reverse-complement ATs, the locus also has runs of pairwise reverse-complementary GTs and ACs (red shading). Presence of such pairwise reverse-complementary VNTRs in the same locus can be explained by a historical template switching event. The secondary structure was inferred with RNAfold (Lorenz et al. 2011).

**Fig S3.**
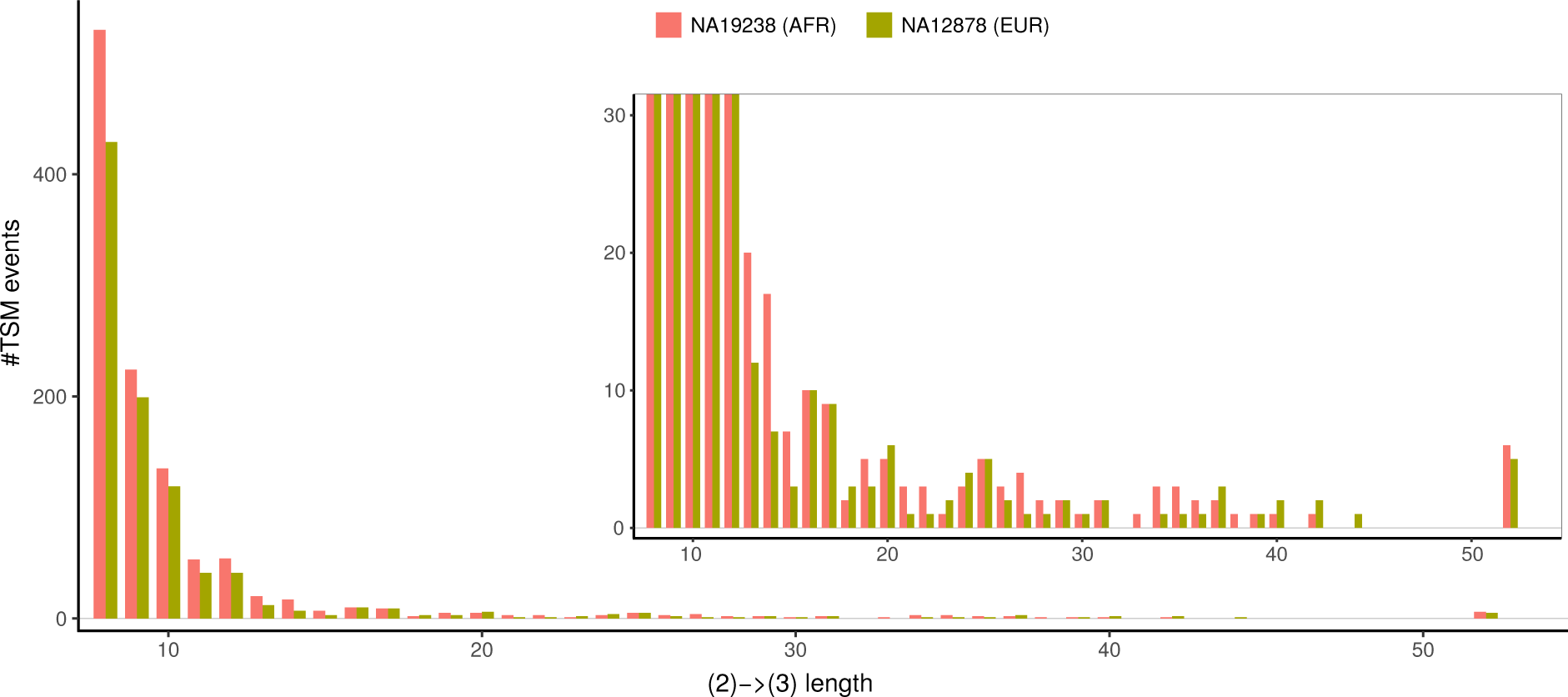
Lengths of inferred TMS events. A great majority of inferred TSM events are 10 bp or shorter in length. Despite the higher numbers of TSM loci identified in African samples, the length distribution of TSM events in individuals of African (NA19238) and European (NA12878) origin are similar. The inset shows lower values. Cases longer than 50 bp are grouped.

**Fig S4.**
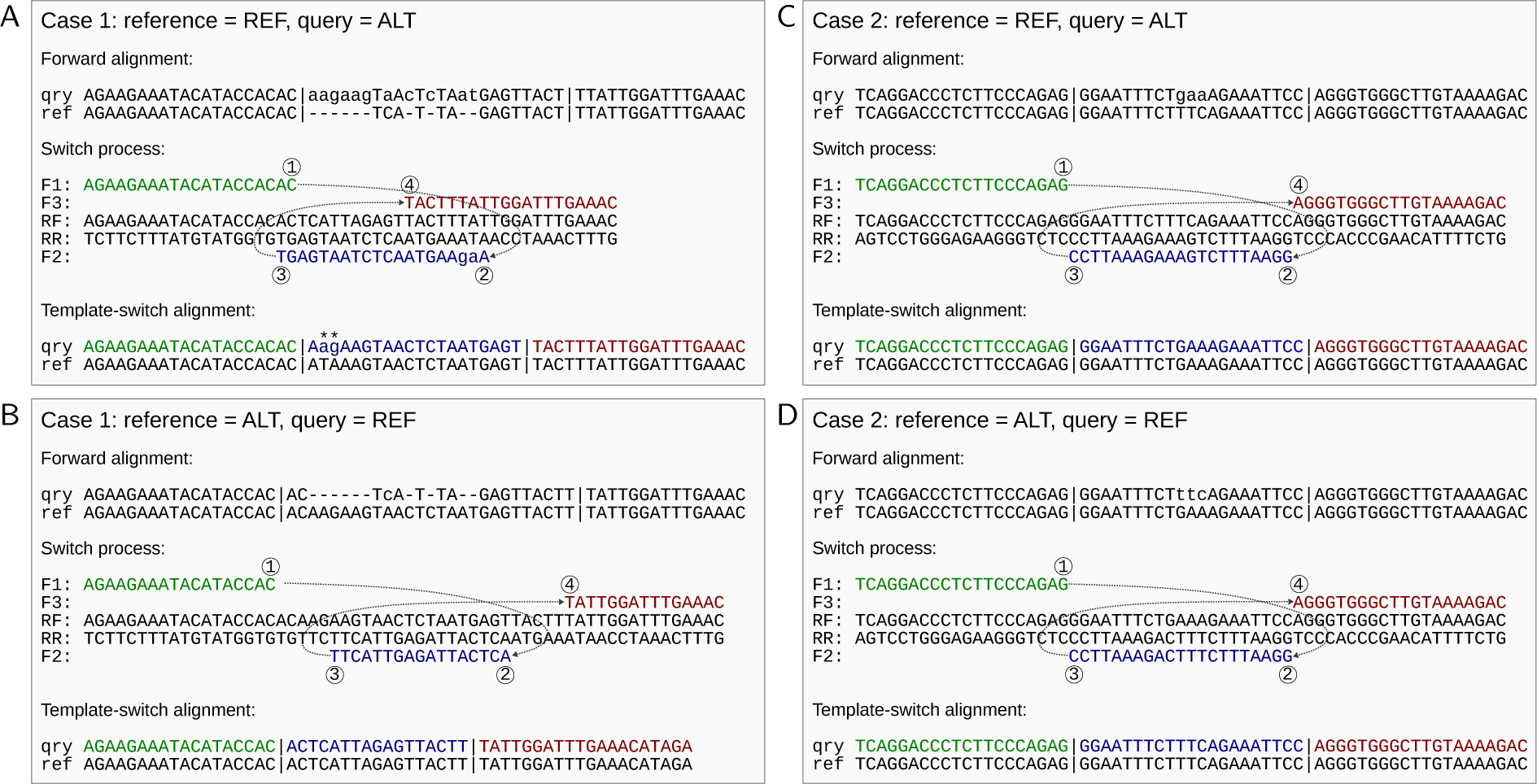
Direction of TSMs. (**A**) The complex mutation (top) is partially explained by the TSM mechanism when the REF allele is used as the reference (middle); the solution (bottom) contains two mismatches (marked with asterisks). (**B**) With the ALT allele as the reference, the complex mutation is fully explained, indicating that ALT is the ancestral state and REF is a derived state. (**C,D**) The inversions in place cannot be polarised as both arrangements fully explain the complex mutation pattern with a single TSM event.

**Fig S5.**
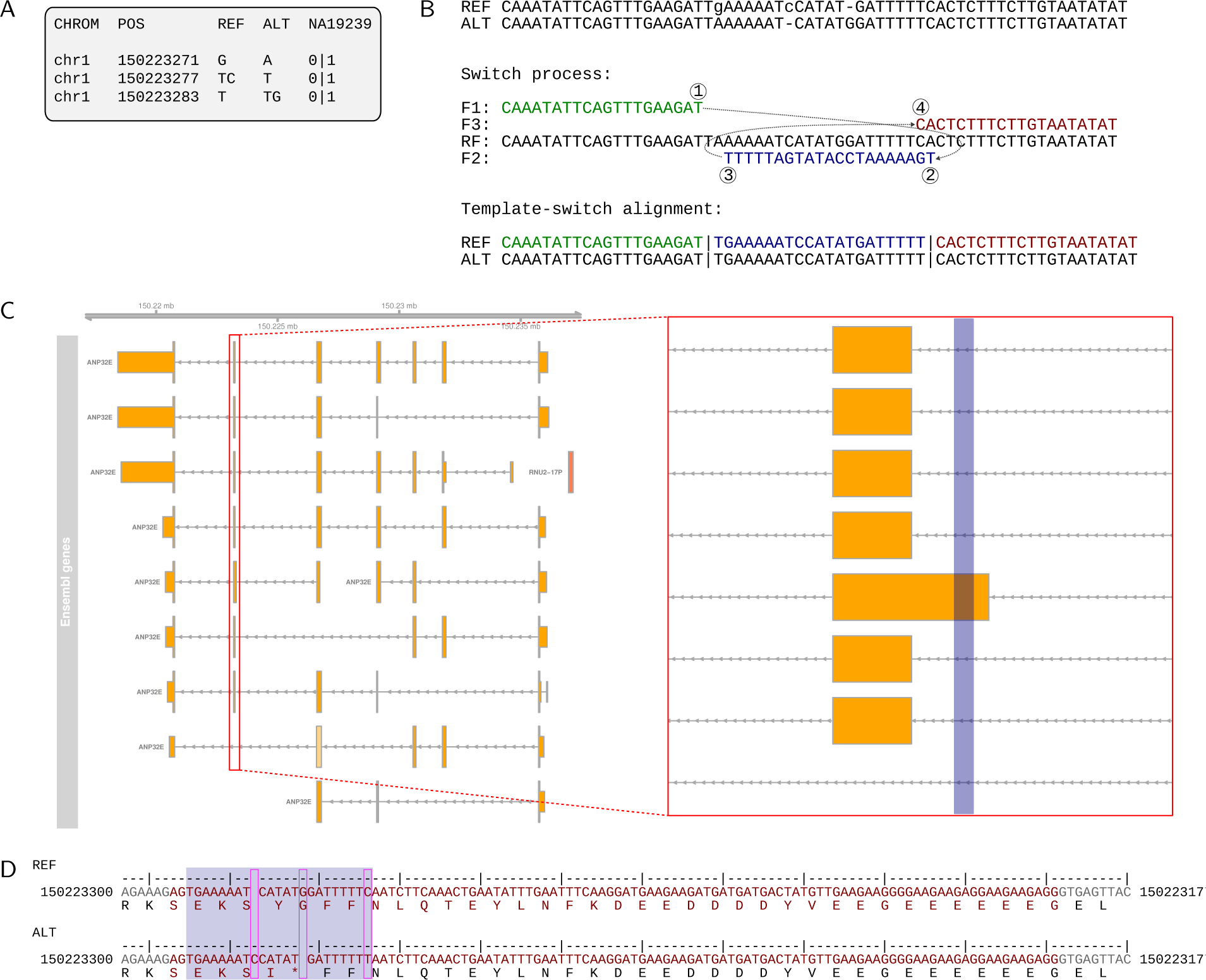
TSM causing an early stop codon. (**A**) The three variants included in the HaplotypeSV data appear together in one individual, NA19239. (**B**) All changes are explained by a template switch event. (**C**) The affected sites are within an alternatively-spliced exon of gene *ANP32E*, highlighted in blue. (**D**) The changes, shown in magenta, alter the reading frame and cause an early stop codon (bottom). Coding sites are shown in red.

**Fig S6.**
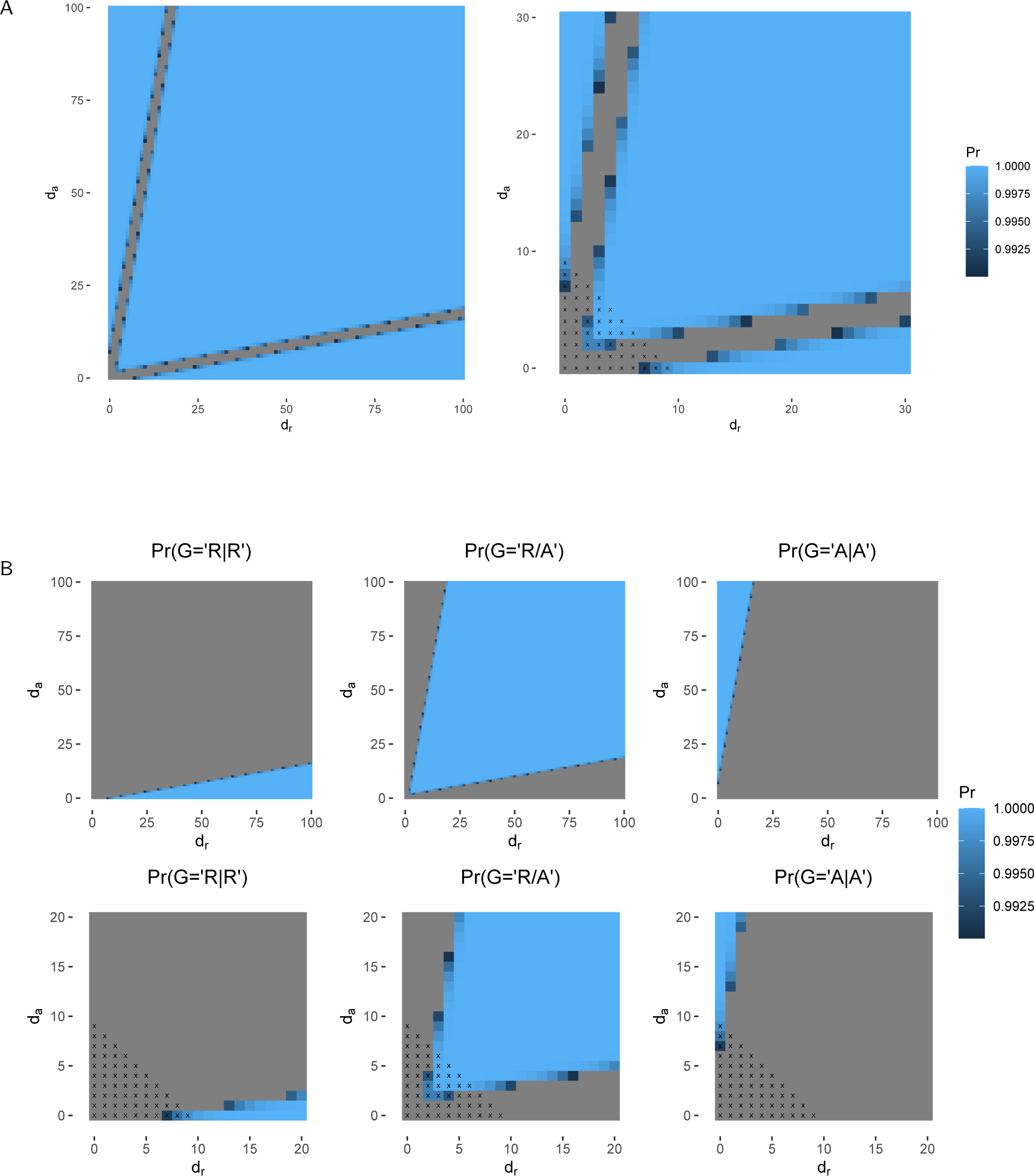
Power of TSM genotyping function. (**A**) The TSM genotype is inferred from mapping coverage of the alternative alleles using a strategy similar to base calling. The heatmaps show the posterior probability of the best genotype for different combinations of REF (*d_r_* ) and ALT (*d_a_*) mapping depths; the combinations indicated with gray have posterior probability *<*0.99 and are considered unknown. The plot on the right shows the values for low depths: the values indicated with crosses are rejected because of low total coverage (*<*10). (**B**) The heatmaps show the posterior probability of the three different genotypes.

**Fig S7.**
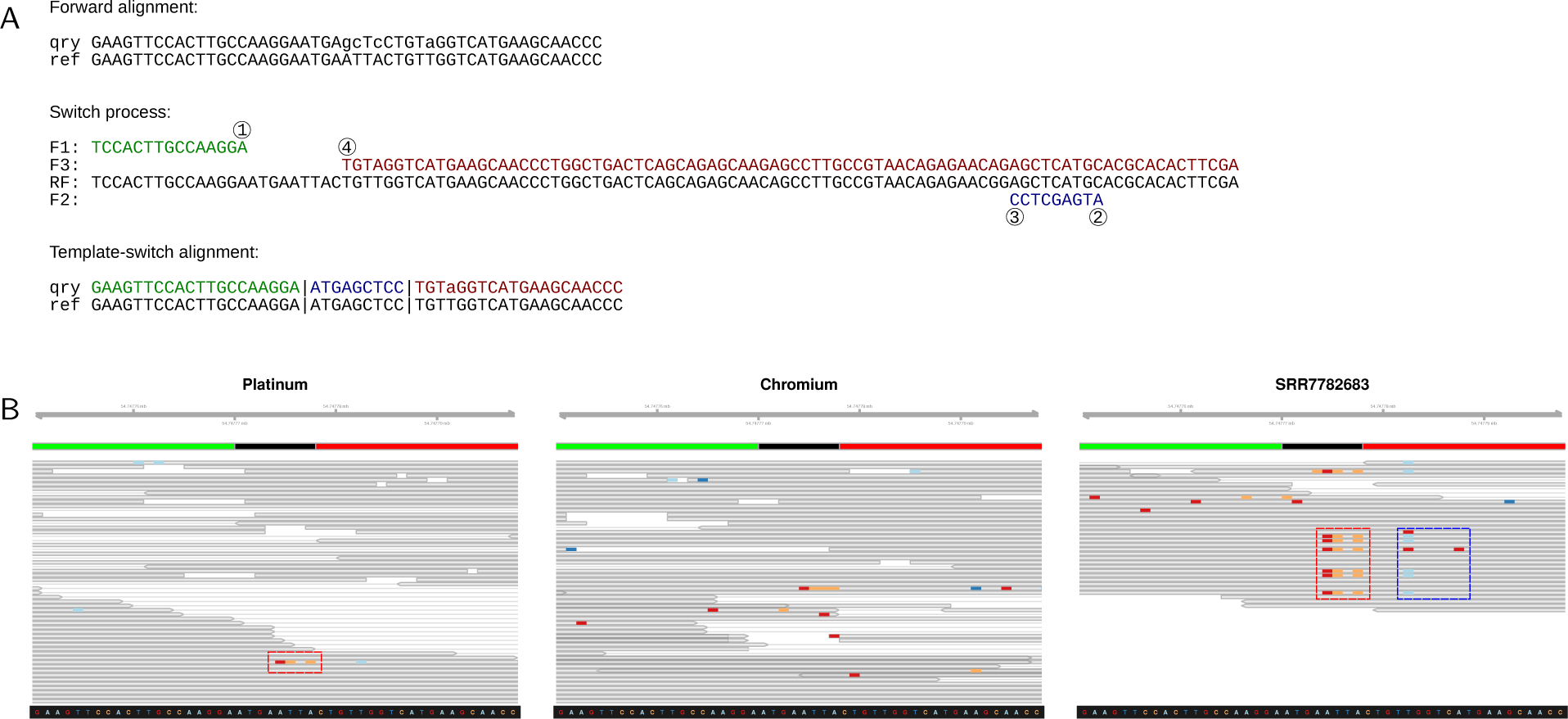
Inconsistency among different short-read datasets. (**A**) The TSM solution explaining a cluster of three base differences (and not explaining one nearby difference). (**B**) The Platinum data have one supporting read (left; red box) whereas the Chromium data have none; an independent dataset has multiple supporting reads (red box), but these are associated with flanking changes (blue box) creating more than two haplotypes. A plausible explanation is the mismapping of reads from another locus. Data were visualized with IGV (Thorvaldsdottir et al. 2013).

**Fig S8.**
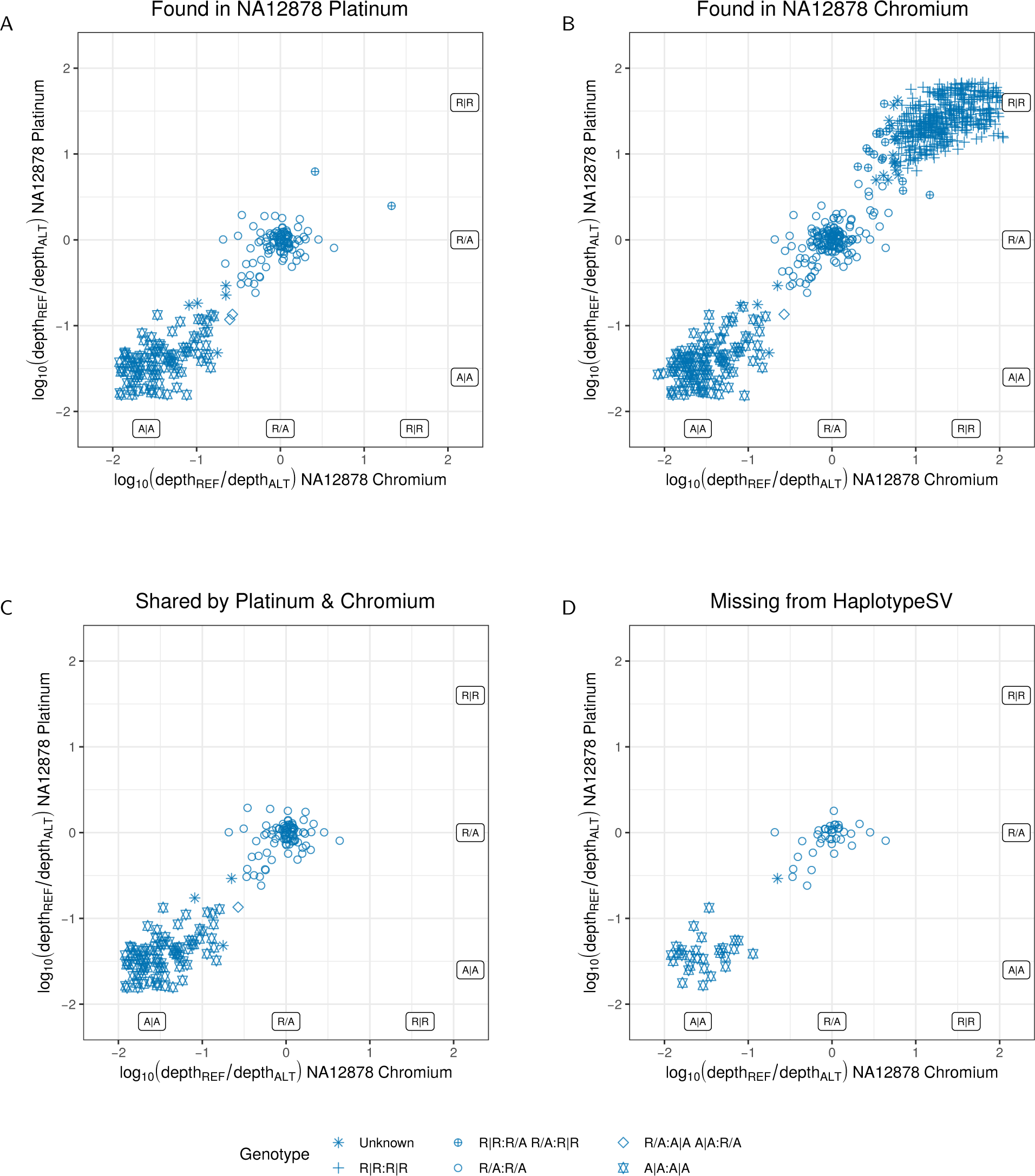
Genotyping of the TSM candidate loci found with short-read data. (**A**) The 211 candidate TSM loci found in NA12878 Platinum data passing the sequence-based filtering. Ratio of REF and ALT allele mapping coverage (*r* = *depth*_REF_*/depth*_ALT_) reflects the genotype: *r ≈* 1 (and thus *log*_10_(*r*) *≈* 0) for a heterozygote; *r «* 1 and *r »* 1 for the two types of homozygotes. The Log10-ratios agree for the NA12878 Platinum and Chromium datasets, and 105 and 96 loci (of the total 211) are called heterozygous and homozygous for ALT by both datasets. (**B**) Of the 755 candidate TSM loci found in NA12878 Chromium data, 438 are called homozygous for REF by both methods. (**C**) The 176 loci found in both short-read datasets. (**D**) The 67 loci found in both short-read datasets but not in HaplotypeSV dataset.

**Fig S9.**
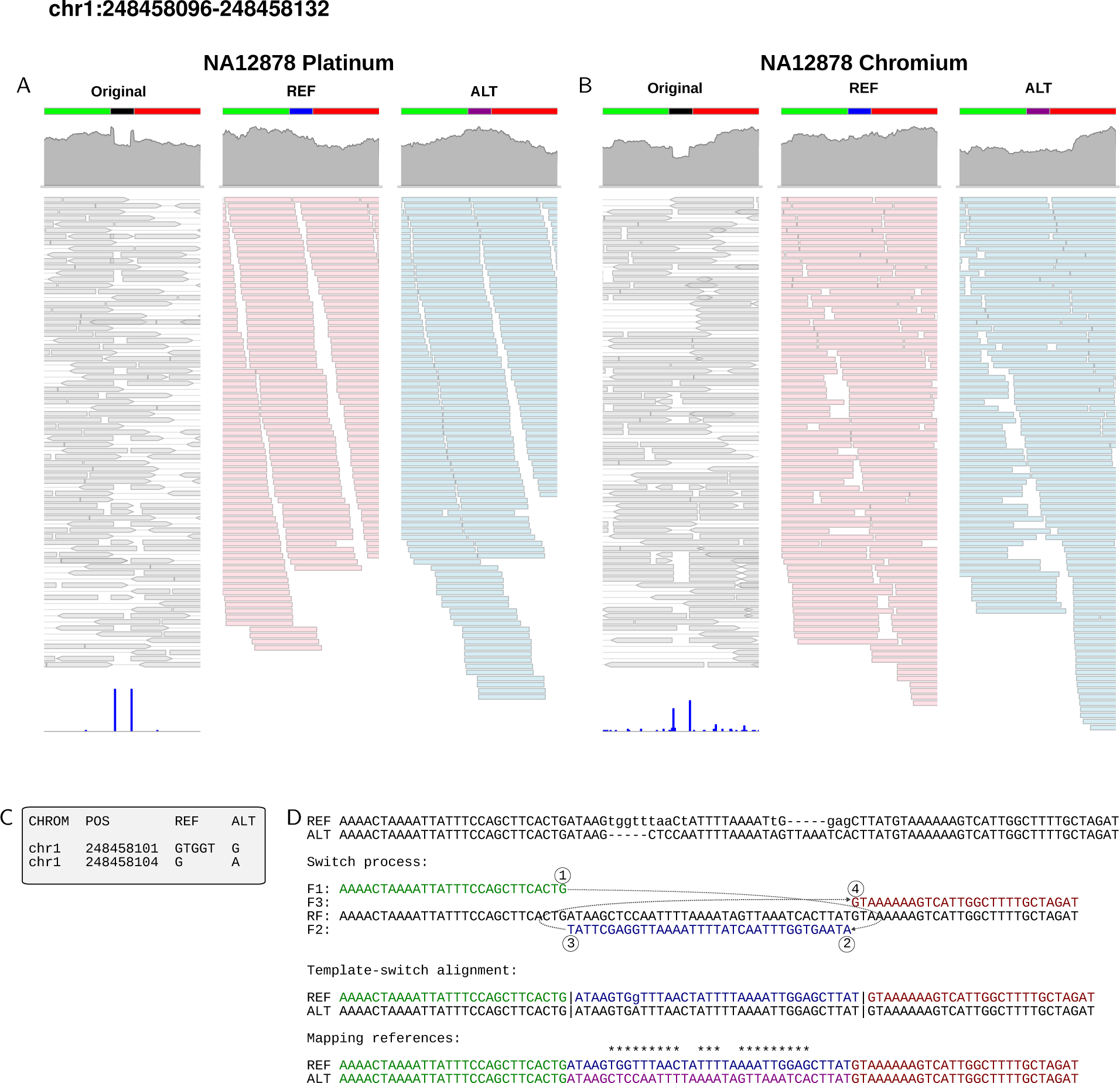
A heterozygous TSM event identified in NA12878 with short-read data. (**A**) Platinum data (left) and (**B**) Chromium data (left) have excess of soft clips (bottom, blue bars) and atypical patterns in the mapping coverage (top, in gray). (**C**) Two variants were called in this region by Ebert et al. (2021) but these fail to explain the haplotype differences. (**D**) De novo assembly of the reads creates two locally highly dissimilar haplotypes (top). All but one difference can be explained with a TSM event, an inversion in place (middle). Using two haplotypes with alternative central parts (blue and magenta; bottom) as the reference, extracted reads map in full length (**A,B**; middle, right) with roughly even coverages. Data were visualised with Gviz (Hahne and Ivanek 2016).

**Fig S10.**
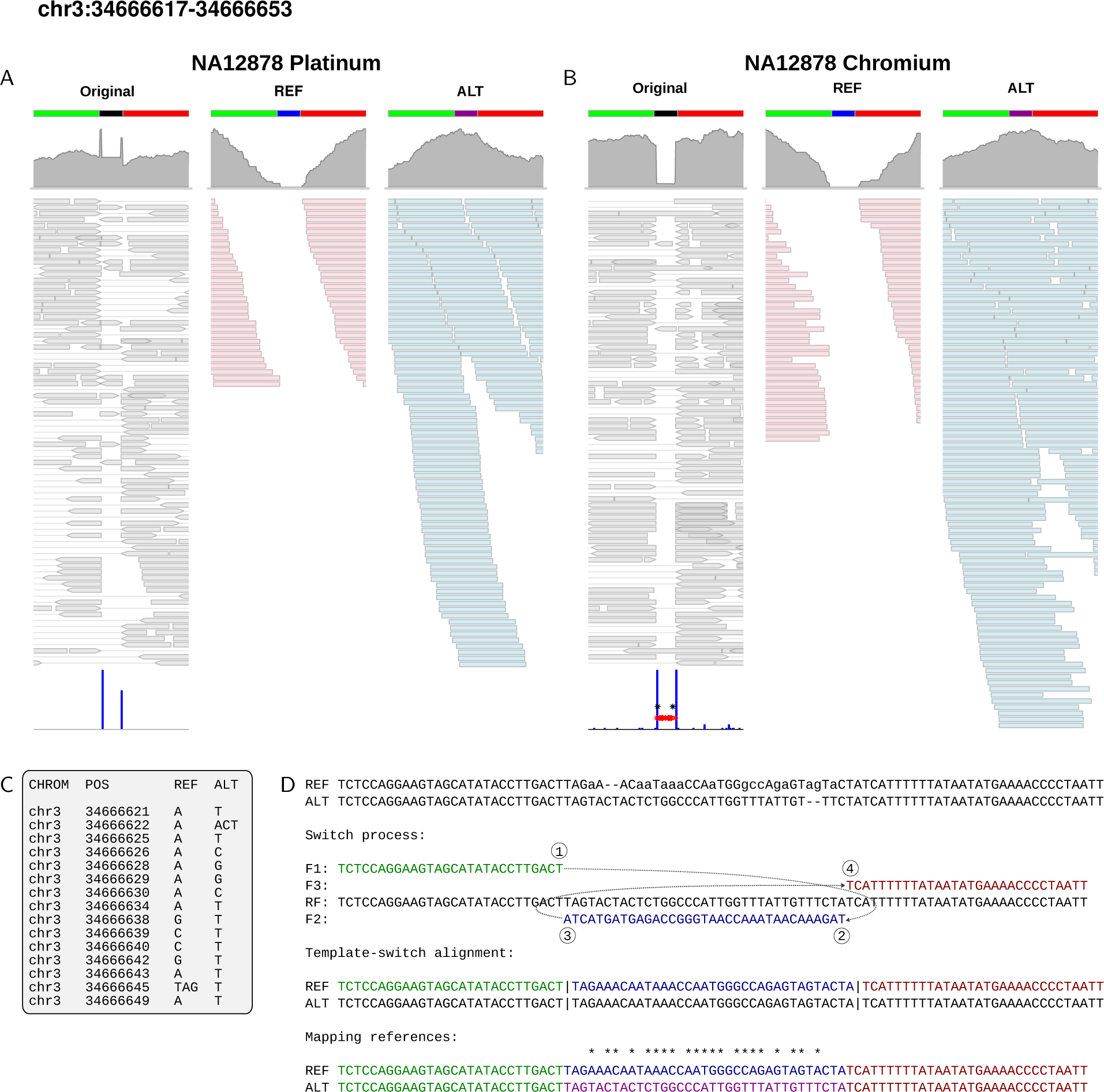
A homozygous TSM event identified in NA12878 with short-read data. (**A**) Platinum data (left) and (**B**) Chromium data (left) have excess of soft clips (bottom, blue bars) and atypical patterns in the mapping coverage (top, in gray). (**C**) Fifteen variants fully explaining the differences were called in the Chromium data while no variants were indicated in this region in the Platinum data or by Ebert et al. (**D**) De novo assembly of the reads creates two locally highly dissimilar haplotypes (top). All differences can be explained with a TSM event, an inversion in place (middle). Using two haplotypes with alternative central parts (blue and magenta; bottom) as the reference, extracted reads map in full length on the ALT haplotype only (**A,B**; middle, right). Data were visualised with Gviz (Hahne and Ivanek 2016).

**Fig S11.**
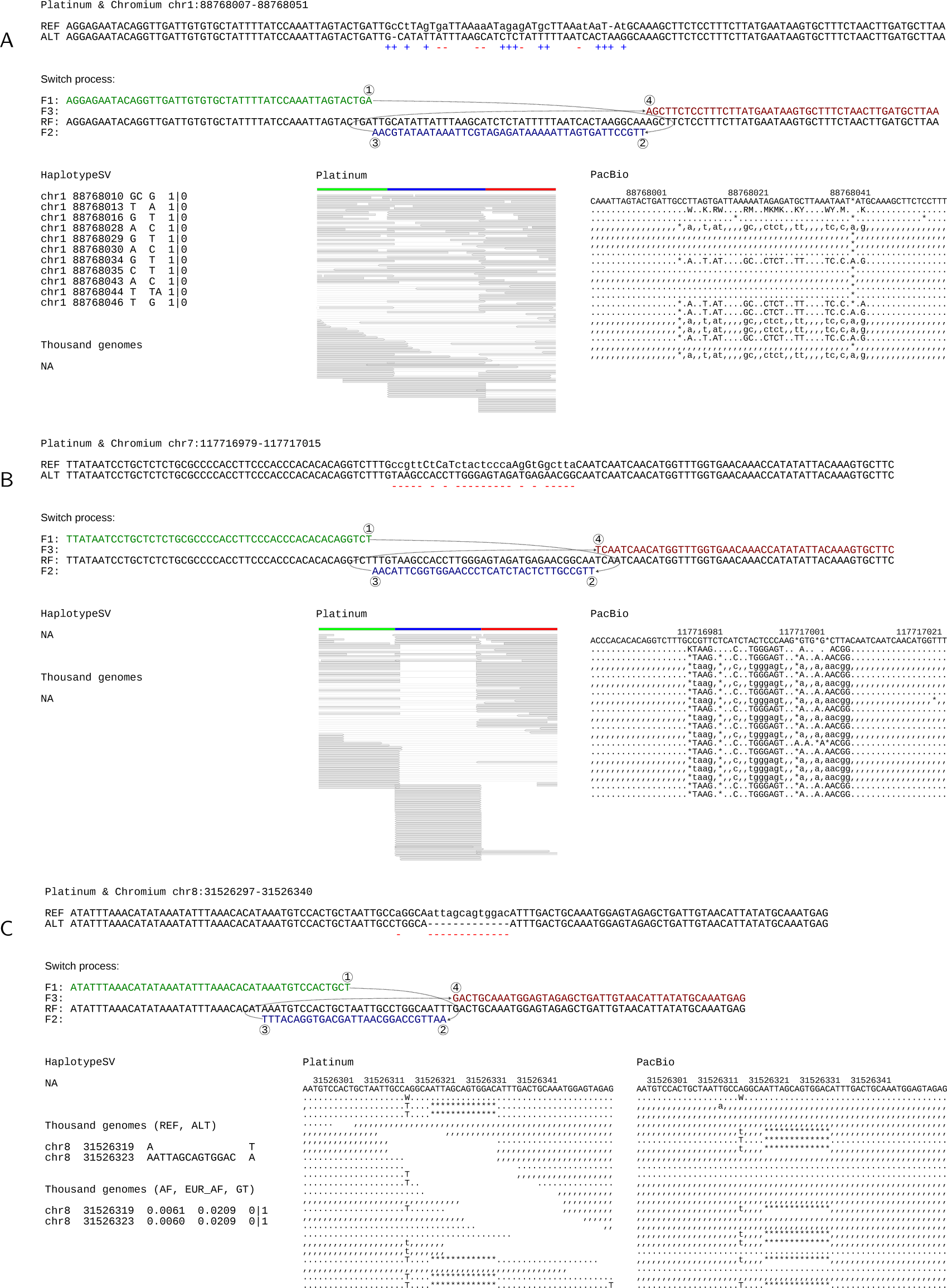
TSM candidate loci missing from HaplotypeSV data. (**A**) A partially called locus. Called and missing changes are indicated with plus (blue) and minus (red) signs. (**B**) A locus completely missing from HaplotypeSV data. (**C**) A locus that is missing despite being in full in the 1000 Genomes (1kG) data. The plots show the two alleles and their TSM solution (top); HaplotypeSV and 1kG variants (left); NA12878 Platinum (middle) and PacBio (right) data for the given locus. 1kG allele frequencies are shown for all and European samples. Alignment gaps cannot be visualised with the Gviz package and in (C) the Platinum data are shown in text format.

**Fig S12.**
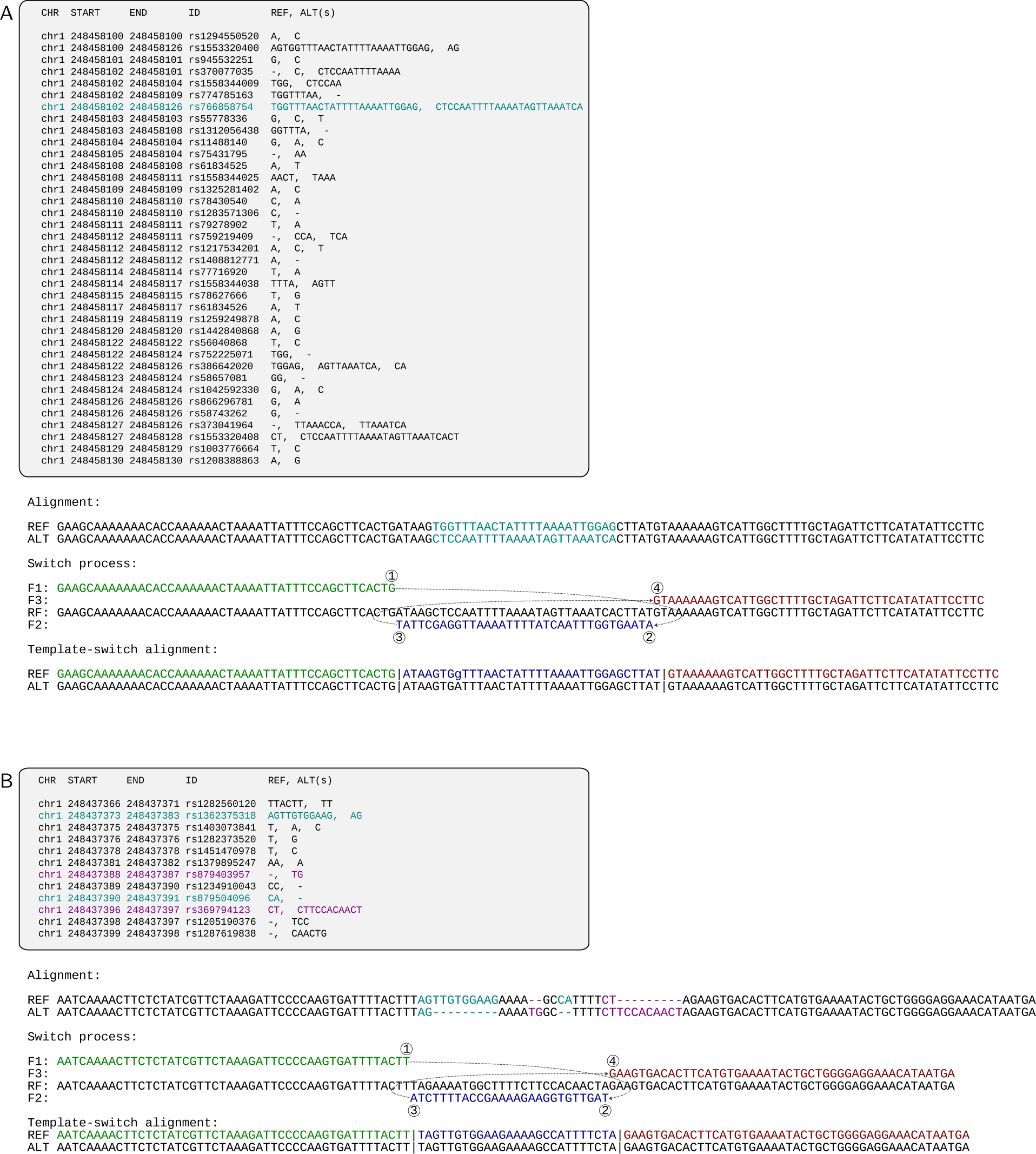
Description of TSM variants in the dbSNP database. (**A**) The TSM locus is described in full with a 25 bp MNV (cyan), but the same changes are also given as multiple SNVs and indels. (**B**) TSM haplotypes can be reconstructed from four different variants (cyan, magenta) but the region has also additional variants annotated.

**Fig S13.**
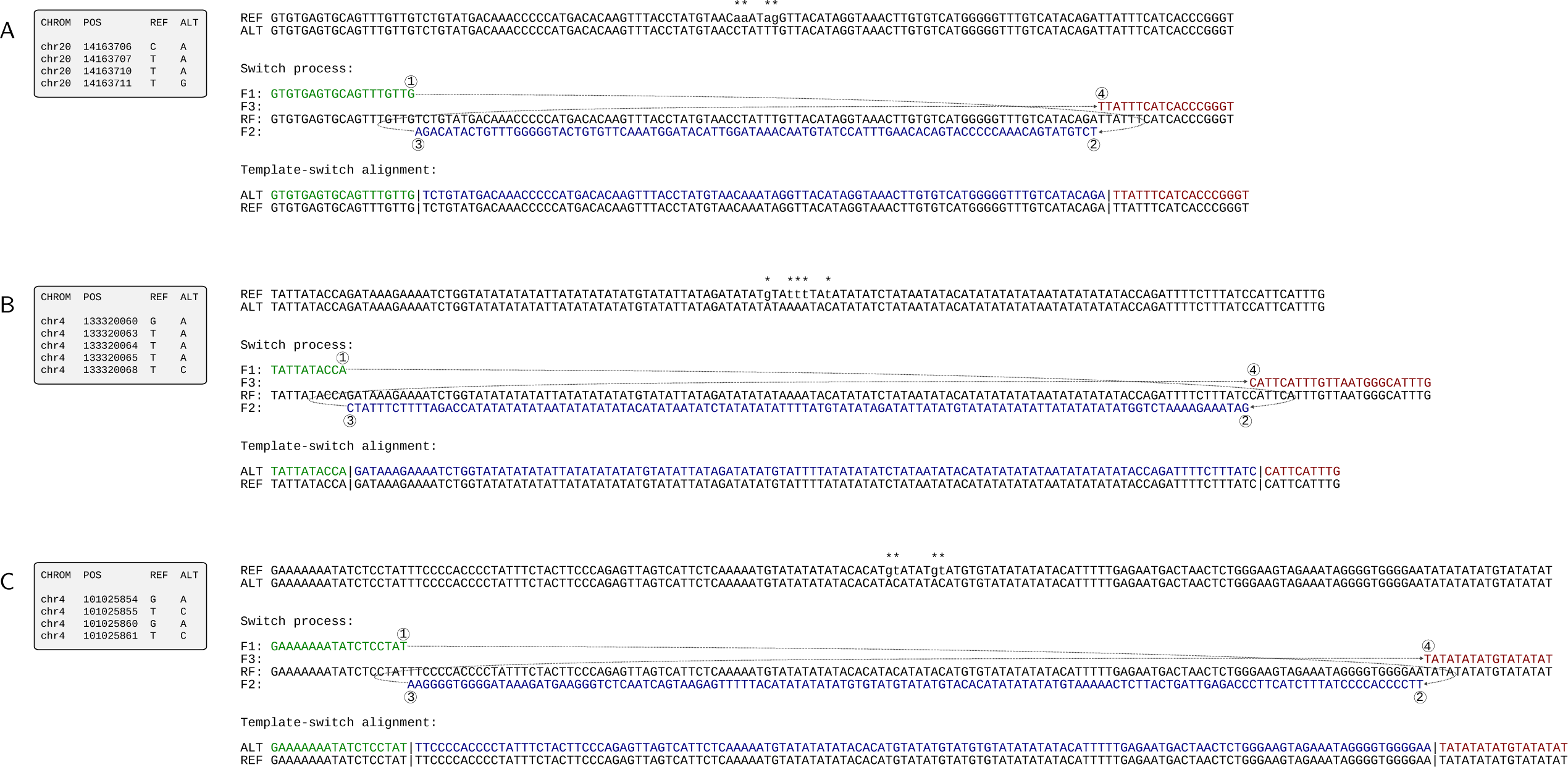
The three longest TSM events explaining gnomAD MNVs. (**A**) The longest TSM event within a complex sequence is 90 bp long and explains an MNV of four base differences. (**B,C**) The longest TSM events involving VNTRs, 119 and 125 bp in length, explain MNVs of five and four base differences.

**Fig S14.**
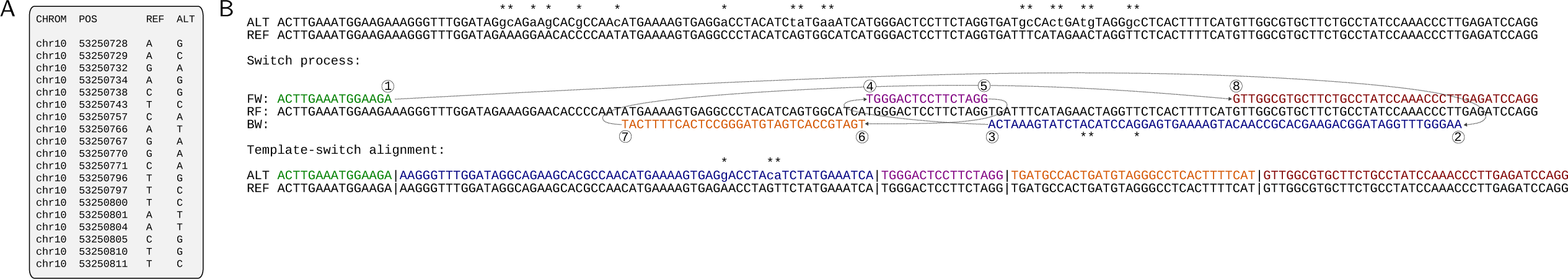
Large gnomAD MNV explained by two TSMs. (**A**) A cluster of 19 base changes in gnomAD MNV data. (**B**) Of the 19 base differences (top), 16 can be explained with two TSM events (middle, bottom). DNA replication proceeds from left to right (green); at ①, the replication jumps to ② on the opposite strand and proceeds in a reverse-complement manner until ③ (blue); it then jumps to ④ in the original template and proceeds until ⑤ (magenta); the replication again changes the template (either to the opposite strand or to the newly synthesized strand), jumps to ⑥ and continues until in reverse-complement manner until (J) (orange); the replication makes a final jump to ® in the original template and continues normally (red).

**Fig S15.**
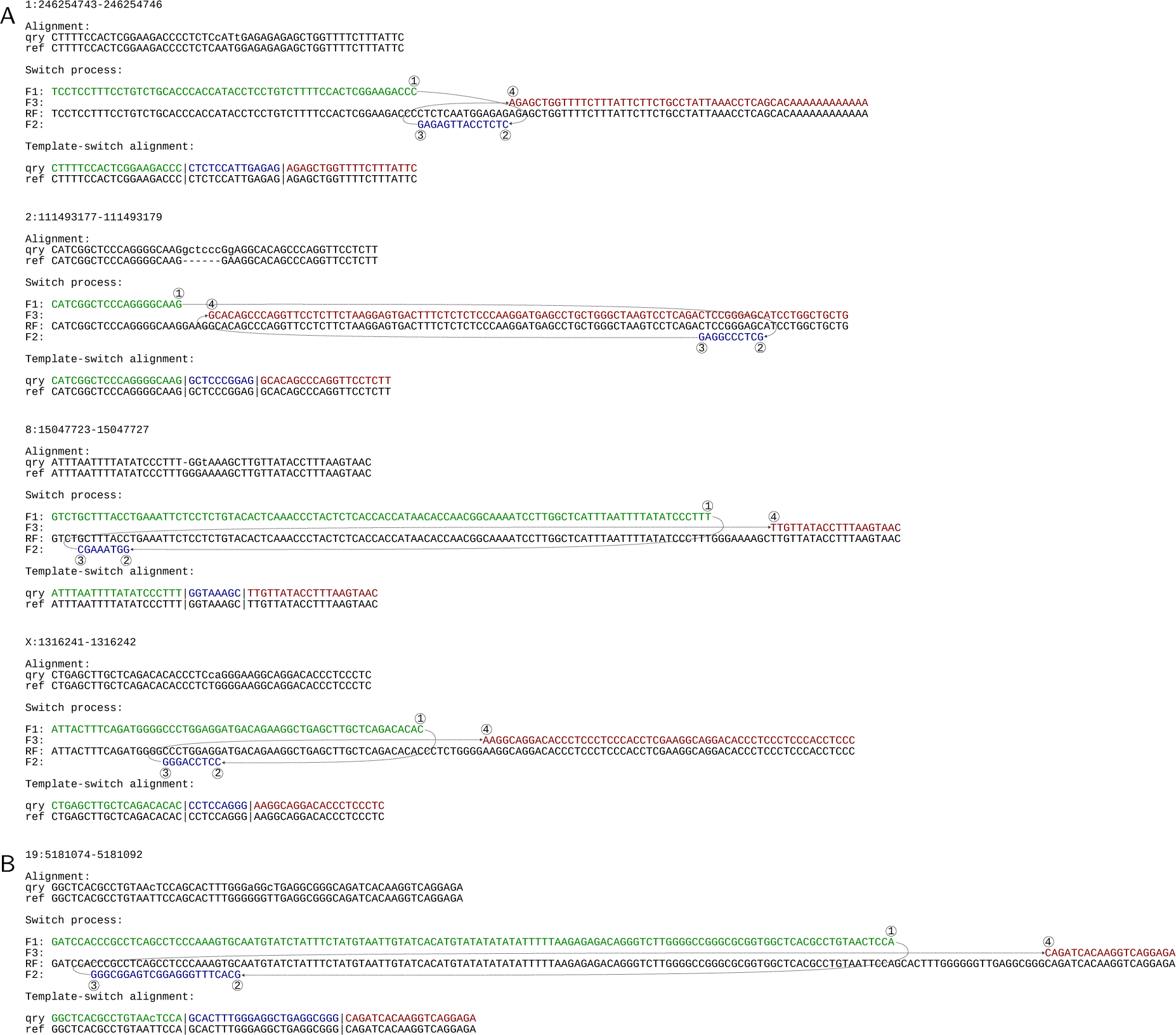
De novo MNVs explainable with the TSM mechanism. (**A**) Four MNVs in the third-generation individuals of Sasani et al. (2019) that can be explained with a single TSM. (**B**) A TSM solution that can explain two of the three clustered de novo SNVs in a third-generation individual of Sasani et al.

**Table S1.** Number and length of inferred TSM events, as well as proportion of homozygous loci and resolved and ancestral (of the former) allelic state in different samples. Only unmasked loci are included.

| ID | Population | Region | n | Length (bp) |  | Allele (%) |  | Polarization (%) |  |
| --- | --- | --- | --- | --- | --- | --- | --- | --- | --- |
|  |  |  |  | average | maxim. | homozyg. | singleton | resolved | ancestral |
| HG02011 | ACB | AFR | 1046 | 10.67 | 171 | 30.50 | 7.58 | 89.50 | 49.68 |
| NA19983 | ASW | AFR | 1077 | 10.77 | 171 | 27.48 | 8.41 | 88.03 | 53.74 |
| HG03125 | ESN | AFR | 1073 | 10.43 | 171 | 29.45 | 7.12 | 88.01 | 51.32 |
| HG03371 | ESN | AFR | 1070 | 10.44 | 171 | 31.40 | 8.43 | 89.09 | 52.57 |
| HG02587 | GWD | AFR | 1097 | 10.81 | 180 | 26.71 | 9.59 | 89.44 | 51.73 |
| HG02818 | GWD | AFR | 1038 | 10.64 | 100 | 29.38 | 8.07 | 88.07 | 53.01 |
| HG03065 | MSL | AFR | 1060 | 10.48 | 171 | 30.94 | 8.81 | 89.84 | 53.09 |
| HG03486 | MSL | AFR | 1064 | 10.90 | 180 | 28.85 | 8.39 | 88.17 | 52.08 |
| NA19238 | YRI | AFR | 1121 | 10.44 | 97 | 28.37 | 8.60 | 90.29 | 51.28 |
| NA19239 | YRI | AFR | 1092 | 10.47 | 171 | 29.30 | 8.91 | 89.03 | 52.46 |
| NA12329 | CEU | EUR | 922 | 10.70 | 180 | 35.03 | 3.07 | 88.84 | 52.32 |
| NA12878 | CEU | EUR | 921 | 10.49 | 171 | 36.48 | 2.61 | 89.48 | 54.42 |
| HG00171 | FIN | EUR | 883 | 10.28 | 171 | 34.65 | 3.63 | 90.05 | 54.27 |
| HG00096 | GBR | EUR | 937 | 10.67 | 171 | 36.07 | 3.11 | 89.23 | 53.17 |
| HG01505 | IBS | EUR | 920 | 10.79 | 180 | 32.83 | 4.03 | 89.46 | 52.61 |
| NA20509 | TSI | EUR | 882 | 10.47 | 180 | 35.49 | 4.52 | 89.12 | 55.34 |
| NA24385 | AZK | EUR | 872 | 10.48 | 171 | 36.24 | 2.71 | 89.58 | 54.86 |
| HG00864 | CDX | EAS | 942 | 10.61 | 171 | 39.17 | 4.33 | 88.67 | 50.42 |
| NA18534 | CHB | EAS | 923 | 10.59 | 171 | 39.98 | 4.34 | 87.99 | 52.15 |
| HG00512 | CHS | EAS | 917 | 10.52 | 171 | 39.91 | 2.82 | 89.12 | 50.67 |
| HG00513 | CHS | EAS | 929 | 10.69 | 180 | 41.87 | 3.57 | 88.41 | 50.24 |
| NA18939 | JPT | EAS | 927 | 10.50 | 171 | 42.29 | 3.16 | 89.76 | 50.30 |
| HG01596 | KHV | EAS | 910 | 10.78 | 171 | 39.01 | 4.12 | 88.50 | 50.00 |
| HG03009 | BEB | SAS | 878 | 10.45 | 171 | 35.76 | 3.85 | 88.52 | 52.76 |
| NA20847 | GIH | SAS | 938 | 10.55 | 171 | 35.82 | 4.59 | 89.15 | 53.46 |
| HG03732 | ITU | SAS | 920 | 10.37 | 171 | 35.87 | 3.57 | 88.15 | 52.77 |
| HG02492 | PJL | SAS | 879 | 10.80 | 180 | 39.59 | 3.14 | 87.94 | 52.39 |
| HG03683 | STU | SAS | 939 | 10.22 | 97 | 33.97 | 4.06 | 89.80 | 50.89 |
| HG01114 | CLM | AMR | 923 | 10.64 | 171 | 33.69 | 3.86 | 87.66 | 53.83 |
| NA19650 | MXL | AMR | 906 | 10.58 | 171 | 35.76 | 3.43 | 88.41 | 52.93 |
| HG00731 | PUR | AMR | 949 | 10.63 | 171 | 34.56 | 3.09 | 88.55 | 52.91 |
| HG00732 | PUR | AMR | 955 | 10.60 | 171 | 34.87 | 3.23 | 88.20 | 51.36 |

**Table S2.** Annotation of the 2,200 TSM candidate loci identified in HaplotypeSV data. A locus can contribute to multiple categories but to each category only once.

| Genomic region | Number of loci |
| --- | --- |
| intergenic | 776 |
| intron | 1342 |
| promoter | 220 |
| threeUTR | 49 |
| fiveUTR | 17 |
| spliceSite | 11 |
| coding | 24 |
| total | 2439 |

**Table S3.** Genotypes inferred with different datasets for variable TSM loci in NA12878 HaplotypeSV data.

| HaplotypeSV | Chromium | Platinum |  |  |  |
| --- | --- | --- | --- | --- | --- |
|  |  | R\|R | R/A | A\|A | NA |
| NA | R\|R | 2 | 0 | 0 | 0 |
| NA | R/A | 0 | 71 | 1 | 3 |
| NA | A\|A | 0 | 2 | 4 | 0 |
| NA | NA | 0 | 0 | 0 | 1 |
| R/A | R\|R | 6 | 1 | 0 | 0 |
| R/A | R/A | 5 | 425 | 0 | 0 |
| R/A | A\|A | 0 | 0 | 2 | 0 |
| R/A | NA | 1 | 1 | 0 | 0 |
| A\|A | R\|R | 2 | 0 | 0 | 0 |
| A\|A | R/A | 0 | 9 | 7 | 0 |
| A\|A | A\|A | 0 | 3 | 310 | 3 |
| A\|A | NA | 0 | 1 | 4 | 0 |

**Table S4.** Parental genotypes for HaplotypeSV TSM loci confirmed variable in NA12878 Chromium data.

| NA12878 | NA12891 | NA12892 |  |  |  |
| --- | --- | --- | --- | --- | --- |
|  |  | R\|R | R/A | A\|A | NA |
| R/A | R\|R | 1 | 80 | 35 | 4 |
| R/A | R/A | 65 | 113 | 53 | 14 |
| R/A | A\|A | 39 | 42 | 2 | 2 |
| R/A | NA | 1 | 20 | 3 | 2 |
| A\|A | R\|R | 0 | 0 | 0 | 0 |
| A\|A | R/A | 0 | 42 | 46 | 1 |
| A\|A | A\|A | 0 | 43 | 159 | 4 |
| A\|A | NA | 0 | 3 | 6 | 3 |

## References

1. Aganezov S et al. 2022. A complete reference genome improves analysis of human genetic variation. Science. 376: eabl3533.

2. Benson G. 1999. Tandem repeats finder: a program to analyze DNA sequences. Nucleic Acids Res. 27: 573–580.

3. Bentley DR et al. 2008. Accurate whole human genome sequencing using reversible terminator chemistry. Nature. 456: 53–59.

4. Besenbacher S et al. 2016. Multi-nucleotide de novo mutations in humans. PLoS Genet. 12: e1006315.

5. Caballero J, Smit AFA, Hood L, and Glusman G. 2014. Realistic artificial DNA sequences as negative controls for computational genomics. Nucleic Acids Res. 42: e99.

6. Chen YH, Keegan S, Kahli M, Tonzi P, Feny o D, Huang TT, and Smith DJ. 2019. Transcription shapes DNA replication initiation and termination in human cells. Nat. Struct. Mol. Biol. 26: 67–77.

7. Choi Y, Chan AP, Kirkness E, Telenti A, and Schork NJ. 2018. Comparison of phasing strategies for whole human genomes. PLoS Genet. 14: e1007308.

8. Conrad DF et al. 2011. Variation in genome-wide mutation rates within and between human families. Nat. Genet. 43: 712–714.

9. Danecek P et al. 2021. Twelve years of SAMtools and BCFtools. GigaScience. 10: giab008.

10. Dutra BE and Lovett ST. 2006. Cis and trans-acting effects on a mutational hotspot involving a replication template switch. J. Mol. Biol. 356: 300–311.

11. Eberle MA et al. 2017. A reference data set of 5.4 million phased human variants validated by genetic inheritance from sequencing a three-generation 17-member pedigree. Genome Res. 27: 157–164.

12. Ebert P et al. 2021. Haplotype-resolved diverse human genomes and integrated analysis of structural variation. Science. 372: eabf7117.

13. Eid J et al. 2009. Real-time DNA sequencing from single polymerase molecules. Science. 323: 133–138.

14. Falconer E and Lansdorp PM. 2013. Strand-seq: a unifying tool for studies of chromosome segregation. Semin. Cell Dev. Biol. 24: 643–652.

15. Hahne F and Ivanek R 2016. Visualizing Genomic Data Using Gviz and Bioconductor. In: Statistical Genomics: Methods and Protocols. Ed. by E Mathe and S Davis. New York, NY: Springer New York, pp. 335–351.

16. Hamperl S, Bocek MJ, Saldivar JC, Swigut T, and Cimprich KA. 2017. Transcription-replication conflict orientation modulates R-loop levels and activates distinct DNA damage responses. Cell. 170: 774– 786.e19.

17. Harris K and Nielsen R. 2014. Error-prone polymerase activity causes multinucleotide mutations in humans. Genome Res. 24: 1445–1454.

18. Hastings PJ, Ira G, and Lupski JR. 2009. A microhomology-mediated break-induced replication model for the origin of human copy number variation. PLoS Genet. 5: e1000327.

19. Holland AJ and Cleveland DW. 2012. Chromoanagenesis and cancer: mechanisms and consequences of localized, complex chromosomal rearrangements. Nat. Med. 18: 1630–1638.

20. International Human Genome Sequencing Consortium. 2004. Finishing the euchromatic sequence of the human genome. Nature. 431: 931–945.

21. Karczewski KJ et al. 2020. The mutational constraint spectrum quantified from variation in 141,456 humans. Nature. 581: 434–443.

22. Lander ES et al. 2001. Initial sequencing and analysis of the human genome. Nature. 409: 860–921.

23. Lawrence M, Huber W, Pages H, Aboyoun P, Carlson M, Gentleman R, Morgan MT, and Carey VJ. 2013. Software for computing and annotating genomic ranges. PLoS Comput. Biol. 9: e1003118.

24. Lee JA, Carvalho CMB, and Lupski JR. 2007. A DNA replication mechanism for generating nonrecurrent rearrangements associated with genomic disorders. Cell. 131: 1235–1247.

25. Levy S et al. 2007. The diploid genome sequence of an individual human. PLoS Biol. 5: e254.

26. Li H. 2013. Aligning sequence reads, clone sequences and assembly contigs with BWA-MEM. arXiv. 1303.3997.

27. Liu P et al. 2011. Chromosome catastrophes involve replication mechanisms generating complex genomic rearrangements. Cell. 146: 889–903.

28. Lorenz R, Bernhart SH, Höoner Zu Siederdissen C, Tafer H, Flamm C, Stadler PF, and Hofacker IL. 2011. ViennaRNA Package 2.0. Algorithms Mol. Biol. 6: 26.

29. Löytynoja A and Goldman N. 2017. Short template switch events explain mutation clusters in the human genome. Genome Res. 27: 1039–1049.

30. Marks P et al. 2019. Resolving the full spectrum of human genome variation using Linked-Reads. Genome Res. 29: 635–645.

31. Menardi C, Schneider R, Neuschmid-Kaspar F, Klocker H, Hirsch-Kauffmann M, Auer B, and Schweiger M. 1997. Human APRT deficiency: Indication for multiple origins of the most common Caucasian mutation and detection of a novel type of mutation involving intrastrand-templated repair. Hum. Mutat. 10: 251–255.

32. Miga KH and Wang T. 2021. The Need for a Human Pangenome Reference Sequence. Annu. Rev. Genomics Hum. Genet. 22: 81–102.

33. Nurk S et al. 2022. The complete sequence of a human genome. Science. 376: 44–53.

34. Obenchain V, Lawrence M, Carey V, Gogarten S, Shannon P, and Morgan M. 2014. VariantAnnotation: a Bioconductor package for exploration and annotation of genetic variants. Bioinformatics. 30: 2076–2078.

35. Perry GH et al. 2007. Diet and the evolution of human amylase gene copy number variation. Nat. Genet. 39: 1256–1260.

36. Porubsky D et al. 2021. Fully phased human genome assembly without parental data using single-cell strand sequencing and long reads. Nat. Biotechnol. 39: 302–308.

37. Quinlan AR and Hall IM. 2010. BEDTools: a flexible suite of utilities for comparing genomic features. Bioinformatics. 26: 841–842.

38. Ripley LS. 1982. Model for the participation of quasi-palindromic DNA sequences in frameshift mutation. Proc. Natl. Acad. Sci. U. S. A. 79: 4128–4132.

39. Rosche WA, Trinh TQ, and Sinden RR. 1997. Leading strand specific spontaneous mutation corrects a quasipalindrome by an intermolecular strand switch mechanism. J. Mol. Biol. 269: 176–187.

40. Sakamoto Y, Sereewattanawoot S, and Suzuki A. 2020. A new era of long-read sequencing for cancer genomics. J. Hum. Genet. 65: 3–10.

41. Sasani TA, Pedersen BS, Gao Z, Baird L, Przeworski M, Jorde LB, and Quinlan AR. 2019. Large, three-generation human families reveal post-zygotic mosaicism and variability in germline mutation accumulation. Elife. 8: 46922.

42. Śegurel L, Wyman MJ, and Przeworski M. 2014. Determinants of mutation rate variation in the human germline. Annu. Rev. Genomics Hum. Genet. 15: 47–70.

43. Seier T, Padgett DR, Zilberberg G, Sutera Jr VA, Toha N, and Lovett ST. 2011. Insights into mutagenesis using Escherichia coli chromosomal lacZ strains that enable detection of a wide spectrum of mutational events. Genetics. 188: 247–262.

44. Seplyarskiy VB et al. 2021. Population sequencing data reveal a compendium of mutational processes in the human germ line. Science. 373: 1030–1035.

45. Sherry ST, Ward MH, Kholodov M, Baker J, Phan L, Smigielski EM, and Sirotkin K. 2001. dbSNP: the NCBI database of genetic variation. Nucleic Acids Res. 29: 308–311.

46. Taliun D et al. 2021. Sequencing of 53,831 diverse genomes from the NHLBI TOPMed Program. Nature. 590: 290–299.

47. The 1000 Genomes Project Consortium et al. 2015. A global reference for human genetic variation. Nature. 526: 68–74.

48. Thorvaldsdottir H, Robinson JT, and Mesirov JP. 2013. Integrative Genomics Viewer (IGV): high- performance genomics data visualization and exploration. Brief. Bioinform. 14: 178–192.

49. Veltman JA and Brunner HG. 2012. De novo mutations in human genetic disease. Nat. Rev. Genet. 13: 565–575.

50. Venter JC et al. 2001. The sequence of the human genome. Science. 291: 1304–1351.

51. Wala JA et al. 2018. SvABA: genome-wide detection of structural variants and indels by local assembly. Genome Res. 28: 581–591.

52. Walker CR, Scally A, De Maio N, and Goldman N. 2021. Short-range template switching in great ape genomes explored using pair hidden Markov models. PLoS Genet. 17: e1009221.

53. Wang Q et al. 2020. Landscape of multi-nucleotide variants in 125,748 human exomes and 15,708 genomes. Nat. Commun. 11: 2539.

54. Weischenfeldt J, Symmons O, Spitz F, and Korbel JO. 2013. Phenotypic impact of genomic structural variation: insights from and for human disease. Nat. Rev. Genet. 14: 125–138.

55. Weisenfeld NI, Kumar V, Shah P, Church DM, and Jaffe DB. 2017. Direct determination of diploid genome sequences. Genome Res. 27: 757–767.

56. Yan SM, Sherman RM, Taylor DJ, Nair DR, Bortvin AN, Schatz MC, and McCoy RC. 2021. Local adaptation and archaic introgression shape global diversity at human structural variant loci. Elife. 10: e67615.

57. Zhang F, Gu W, Hurles ME, and Lupski JR. 2009. Copy number variation in human health, disease, and evolution. Annu. Rev. Genomics Hum. Genet. 10: 451–481.

58. Zhao H, Sun Z, Wang J, Huang H, Kocher JP, and Wang L. 2014. CrossMap: a versatile tool for coordinate conversion between genome assemblies. Bioinformatics. 30: 1006–1007.

59. Zook JM et al. 2016. Extensive sequencing of seven human genomes to characterize benchmark reference materials. Sci Data. 3: 160025.

